# Size Scaling of the Electrochemical Performance of Ti_3_C_2_T*_x_* MXene Microelectrode Arrays for Electrophysiological Recording and Stimulation

**DOI:** 10.64898/2026.03.19.712102

**Authors:** Spencer R. Averbeck, Raghav Garg, Royce Dong, Daryl Hurwitz, Nicholas V. Apollo, Michael S. Beauchamp, Flavia Vitale

**Affiliations:** S. R. Averbeck, R. Dong, F. Vitale Department of Bioengineering, University of Pennsylvania, Philadelphia, Pennsylvania – 19104, USA; S. R. Averbeck, R. Garg, R. Dong, D. Hurwitz, N.V. Apollo, F. Vitale Center for Neuroengineering and Therapeutics, University of Pennsylvania, Philadelphia, Pennsylvania – 19104, USA; R. Garg, F. Vitale Department of Neurology, University of Pennsylvania, Philadelphia, Pennsylvania – 19104, USA; M. S. Beauchamp Department of Neurosurgery, University of Pennsylvania, Philadelphia, Pennsylvania – 19104, USA; F. Vitale Department of Physical Medicine and Rehabilitation, University of Pennsylvania, Philadelphia, Pennsylvania – 19104, USA; F. Vitale Center for Neurotrauma, Neurodegeneration, and Restoration, Corporal Michael J. Crescenz Veterans Affairs Medical Center, Philadelphia, Pennsylvania – 19104, USA

**Keywords:** Microelectrodes, Implantables, Stimulation, MXene, Ti_3_C_2_T*_x_*, Electrochemistry

## Abstract

Miniaturized neural interfaces for research, diagnostics, and neuromodulation therapies require electrode materials that maintain low impedance and high charge injection capacity as device dimensions shrink to ensure high-quality recordings and safe stimulation. Conventional interfaces rely on metals like platinum (Pt), which are limited by intrinsically high impedance and low charge transfer capacity, reducing their performance in sub-100 µm applications. Ti_3_C_2_T*_x_* MXene has emerged as a promising alternative for high-density recording and stimulation interfaces, though the fundamental charge transfer mechanisms governing its performance remain poorly understood. This study evaluates Ti_3_C_2_T*_x_* MXene microelectrodes across a range of diameters (25 - 500 µm) and systematically elucidates the mechanisms governing their recording and stimulation capabilities. Electrochemical impedance spectroscopy, cyclic voltammetry, and voltage transient measurements – supported by equivalent-circuit modeling – revealed enhanced recording and stimulation capabilities of the MXene microelectrodes over size-matched Pt microelectrodes, attributed to reduced charge-transfer resistance and increased double-layer capacitance. Finally, varying the volume and concentration of the spray-coated Ti_3_C_2_T*_x_* films showed that increased MXene concentration and volume enhanced performance by creating thicker, rougher interfaces. Together, these results establish Ti_3_C_2_T*_x_* MXene as a promising electrode material with exceptional performance at the microscale.

## INTRODUCTION

Bioelectronic devices are increasingly central to modern medicine, enabling both the recording and modulation of electrophysiological activity for diagnostic and therapeutic purposes, respectively. These systems span a wide range of clinical and research applications, interfacing with the heart, muscles, gastrointestinal tract, peripheral nerves, and the central nervous system (CNS). Cardiac pacemakers,^1^ electromyography systems,^2^ vagus nerve stimulators,^3^ and bladder control implants^4^ are just a few examples of technologies that depend on reliable bioelectronic interfaces to sense and respond to complex biological signals. Among these, neural interfaces have emerged as a particularly dynamic area of development, powering devices like cochlear implants, deep brain stimulators (DBS), responsive neurostimulators (RNS), and spinal cord stimulators (SCS) to treat conditions ranging from hearing loss^5^ and Parkinson’s disease^6^ to epilepsy^7^, depression,^8^ and chronic neuropathic pain.^9^ Likewise, brain-computer interfaces (BCIs) are advancing rapidly to restore function and communication in individuals with paralysis or sensory loss.^10,11^ Whether used for recording, stimulation, or both, the performance of bioelectronic technologies depends critically on the quality of signal transduction at the tissue-device interface. As these systems become more sophisticated and miniaturized, there is an increasing need for electrode materials that can reliably and efficiently interface with and control electrically active tissues across a range of physiological size scales. Meeting this need ultimately requires electrode materials that not only integrate mechanically and biologically with tissues but also meet stringent electrochemical performance criteria for both electrophysiological recording and stimulation.

In particular, high-fidelity bioelectronic interfaces for neural recording must maintain low impedance across physiologically relevant frequencies (1 Hz – 1000 Hz), as this reduces thermal noise and improves the signal-to-noise ratio (SNR) in the recordings.^12^ This can most commonly be accomplished either by increasing the electrochemically active surface area through modifying topographic features with pores or increased roughness, or by using materials with low inherent electrochemical impedance due to kinetically fast surface redox reactions.^13^ Likewise, for effective and safe electrical stimulation, electrodes must have high charge injection capacity (CIC) and ideally promote primarily capacitive or pseudocapacitive charge transfer to minimize adverse irreversible faradaic reactions that may be harmful to the tissues and degrade the system over time^13,14^

As demands grow for more targeted high-resolution bioelectronic interfaces with both electrical recording and stimulation capabilities, such as cortical microelectrodes for mapping epileptic foci^15–17^ and high-fidelity brain-computer interfaces for restoring communication,^18–20^ – there is an increasing push toward high-density microelectrode arrays integrating 100s to 1000s of sub-millimeter contacts. Unfortunately, miniaturizing metal electrodes to achieve more targeted recording and stimulation poses significant challenges. Historically, most neural interfaces have used metals such as platinum (Pt) and gold (Au), and while this has been effective, metal electrodes face limitations at smaller scales due to their intrinsically high impedance across local field potential frequencies (LFP, < 300 Hz).^12^ Additionally, metal electrodes pose limits for providing safe electrical microstimulation due to their low CIC (∼150 µC cm^-2^ in the case of platinum) and greater reliance on faradaic charge transfer reactions, which are chronically less biocompatible and can lead to electrode degradation.^21,22^

To address these limitations, alternative electrode materials such as iridium oxide, nanostructured materials, and conductive polymers have been explored. Iridium oxide films and coatings can significantly boost CIC by leveraging reversible redox reactions and a rougher electrochemically active surface area, allowing for smaller electrode contacts while maintaining efficacious electrical stimulation and high SNR recording capabilities.^23,24^ Other emerging materials, including carbon nanotubes,^25,26^ laser-induced porous graphene,^27^ 3D “fuzzy” graphene,^28,29^ reduced graphene oxide,^30,31^ platinum nanorods,^16,32^ and PEDOT: PSS,^33–37^ offer similarly improved performance by increasing the electrochemically active surface area through pores, layers, and other 3D geometries, which enhances capacitive charge transfer at the electrode-electrolyte interface.

Among these, Ti_3_C_2_T*_x_* MXene, a two-dimensional nanomaterial, has emerged as a promising candidate for neural recording and stimulation electrodes.^38–42^ MXenes are transition-metal carbides and nitrides synthesized by selectively etching the A element, such as aluminum, from their MAX-phase precursors, where M is an early transition metal and X is carbon or nitrogen. Ti_3_C_2_T*_x_* MXene, in particular, offers high electrical conductivity (greater than 20,000 S cm^-1^),^43^ substantial volumetric capacitance (1,500 F cm^-3^),^44^ and has been reported to exhibit favorable biocompatibility in many prior studies,^42,45–54^ making it well-suited for bioelectronic applications. Indeed, Ti_3_C_2_T*_x_* MXene has already been successfully integrated into fabrication processes for wearable cardiac (ECG),^55^ muscle (EMG),^56,57^ and brain (EEG) sensors,^55,58^ as well as for invasive electrocorticography (ECoG) and subdural microelectrode arrays and fibers for recording and stimulation.^39–42,59^ Despite these promising applications, however, the underlying mechanisms of charge-transfer at the electrode-electrolyte interface of Ti_3_C_2_T*_x_* electrodes, and their dependence on electrode size and film processing, remain elusive. This knowledge is essential for advancing MXene-based bioelectronic interfaces, especially as electrode diameters shrink to near-neuron dimensions.

Here, we present a systematic investigation of the electrochemical behavior of Ti_3_C_2_T*_x_* MXene thin-film electrodes in the context of electrophysiological recording and stimulation. Specifically, we characterize mechanisms of charge transfer and transduction of thin-film Ti_3_C_2_T*_x_* electrode arrays in simulated physiological media, and report how they vary with contact size, film properties, and fabrication conditions. First, we analyze the electrochemical impedance of the multi-scale microelectrode contacts ranging in diameter from 25 µm to 500 µm. To benchmark and compare the charge transfer behavior of Ti_3_C_2_T*_x_* against state-of-the-art microelectrode materials, we conduct a comparative study with size-matched thin-film sputtered Pt microelectrodes. We then fit the experimental impedance data to an equivalent circuit model to further elucidate the behavior at the electrochemical interface of Ti_3_C_2_T*_x_* and Pt microelectrodes in aqueous electrolytes. Next, we investigate the charge storage capacity, maximum current limits, and CIC at varying pulse widths for both microelectrode materials across all contact sizes. Finally, we evaluate the effects of varying film processing parameters such as Ti_3_C_2_T*_x_* dispersion concentration and spray-coated volume on the electrochemical and charge-transfer performance of the devices.

## RESULTS

### Fabrication and characterization of the Ti_3_C_2_T*_x_* and Pt microelectrode arrays

Ti_3_C_2_T*_x_* thin-film microelectrode arrays were fabricated on machine-grade silicon wafers using standard cleanroom microlithography techniques under previously established protocols (**Figure 1A, B**).^60^ Briefly, the arrays were fabricated on a 3-inch wafer with a 4 µm-thick Parylene-C bottom layer. The microelectrodes were arranged in a 2 x 8 grid (3 mm x 4 mm) with each column containing eight unique contact sizes with diameters (from top to bottom): 500 µm, 350 µm, 200 µm, 150 µm, 100 µm, 75 µm, 50 µm, and 25 µm (**Figure 1C**). Ti_3_C_2_T*_x_* (3 mg mL^-1^, 100 mL) was spray-cast and patterned onto the devices to form the microelectrode contacts (film thickness: 1.07 µm ± 93.20 nm). The completed devices were inspected with optical and scanning electron microscopy (SEM), and atomic force microscopy (AFM) was used to measure the average root-mean-square (RMS) surface roughness of 19.03 nm ± 2.26 (**Figure 1D**). Energy Dispersive X-Ray spectroscopy (EDX) confirmed the elemental composition of Ti_3_C_2_T*_x_* (C, Ti, O, F) and the encapsulating Parylene-C (chemical formula: C_16_H_14_Cl_2_), with atomic percentages consistent with non-oxidized Ti_3_C_2_T_x_.^61^ Additionally, X-Ray Photoelectron spectroscopy (XPS), X-Ray Diffraction spectroscopy (XRD), and Raman spectroscopy confirmed that the bond, lattice, and flake structure of the Ti_3_C_2_T_x_ were comparable to those of properly synthesized Ti_3_C_2_T_x_ MXene (**Figure S1**).^62^ Size-matched Pt microelectrode arrays were fabricated on Parylene-C substrates following a similar process, but the electrode contacts were instead made by sputtering 200 nm of Pt, rather than spray-casting MXene (**Figure S2A**). The RMS surface roughness of platinum microelectrodes measured via AFM was 41.58 nm ± 4.40 nm **(Figure S2B**).

**Figure 1.**
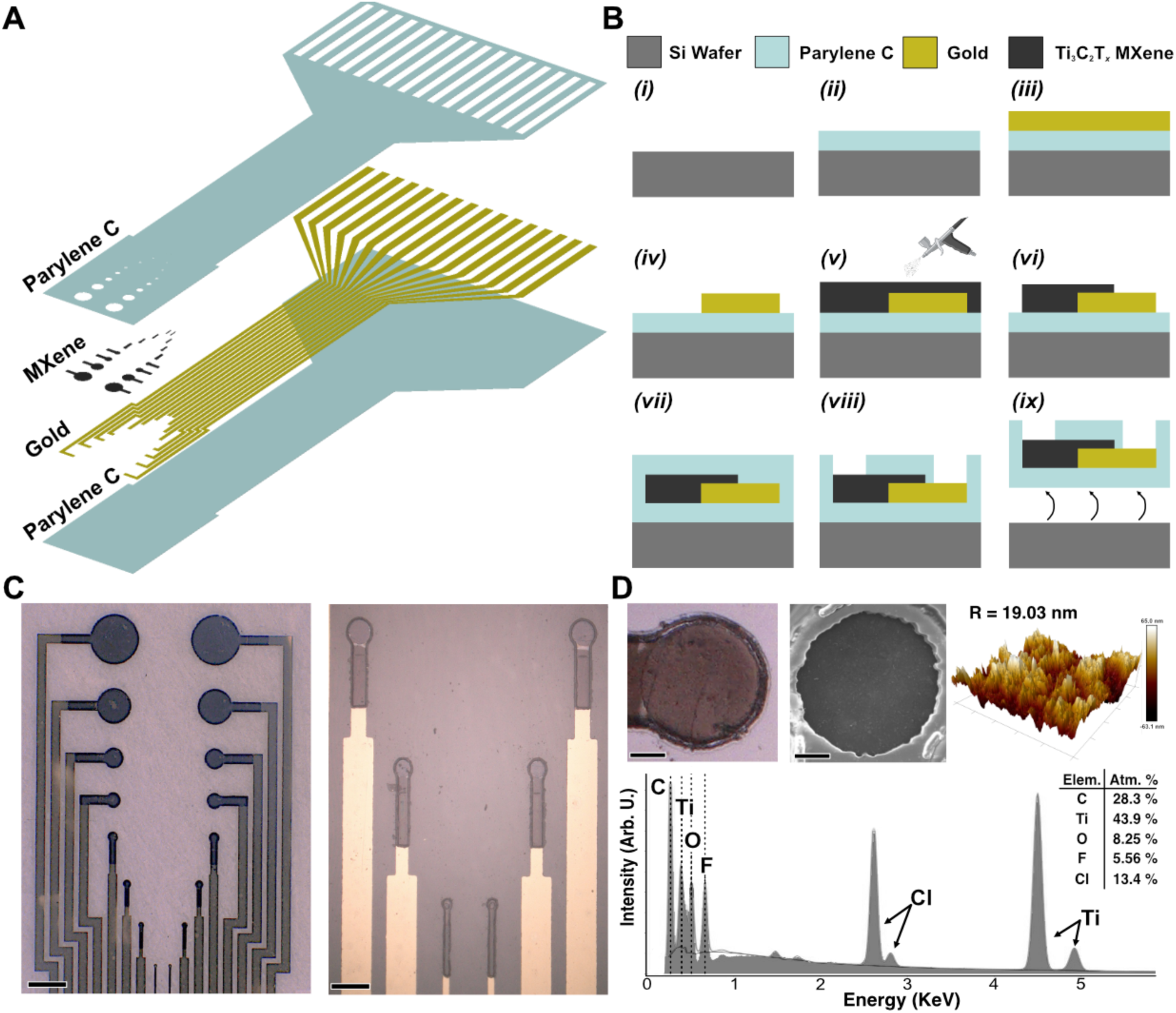
Fabrication and characterization of Ti_3_C_2_T*_x_* microelectrodes. **(A)** Exploded view of the Ti_3_C_2_T*_x_* microelectrode array, with the different material layers. (**B)** Schematic of the microfabrication process. **(C)** *Left:* Digital microscopy image of the Ti_3_C_2_T*_x_* microelectrode array.

Scale bar 500 µm. *Right:* Magnified area highlighting the 75, 50, and 25 µm contacts. Scale bar 150 µm. **(D)** *Top:* (from left to right) optical, SEM, and AFM images of a 100 µm contact (3 mg mL^-1^, 100 mL, film thickness: 1.07 µm ± 93.20 nm). Scale bar 30 µm. *Bottom:* EDX spectra of a Ti_3_C_2_T*_x_* microelectrode.

### Electrochemical impedance spectroscopy

Electrochemical impedance spectroscopy (EIS) of Ti_3_C_2_T*_x_* and Pt microelectrodes was measured in 1x PBS using a standard three-cell configuration with a single-frit Ag/AgCl reference electrode filled with 1 M KCl, a large graphite-rod counter electrode, and a Ti_3_C_2_T*_x_* MXene or Pt microelectrode contact as the working electrode. EIS was acquired across a frequency range of 1 Hz - 100 kHz for all electrode contacts with diameters ranging from 25 µm to 500 µm. (**Figure 2A**). Qualitatively, the phase response of the MXene microelectrodes at all sizes appears to be predominantly resistive, especially around 1 kHz, where the phase of all MXene contact sizes is approximately -15°, with a slow drift towards more capacitive values between -40° and -50° as the frequency decreases to 1 Hz. This near-zero degree resistive phase response across the broadband frequency range (1 kHz – 10 Hz) can be seen manifested in the remarkably stable impedance modulus that maintains a low increase in magnitude with decreasing frequency, averaged across contact sizes as -57.90 ± 5.90 Ω / Hz. In contrast, the Pt microelectrodes show the well-established pseudocapacitive phase across a broad frequency range, where at 1 kHz the phase for all microelectrodes is ∼-70° and swings up to more resistive values ∼-30° as the frequency approaches 1 Hz (**Figure 2B**). The broadband negative shift of the overall phase response thus translates to a steeper slope of the impedance modulus Vs. frequency response curve, calculated at -1,312 ± 159 Ω / Hz. The ∼20x smaller slope of impedance modulus Vs. frequency response and near-resistive phase across all Ti_3_C_2_T*_x_* MXene microelectrodes indicate, in a practical sense, that signal attenuation at the Ti_3_C_2_T*_x_* interface will be minimal across a wide range of frequencies, from action potentials down to local fields and alpha-theta activity, with little to no phase distortion.^12,63^

**Figure 2.**
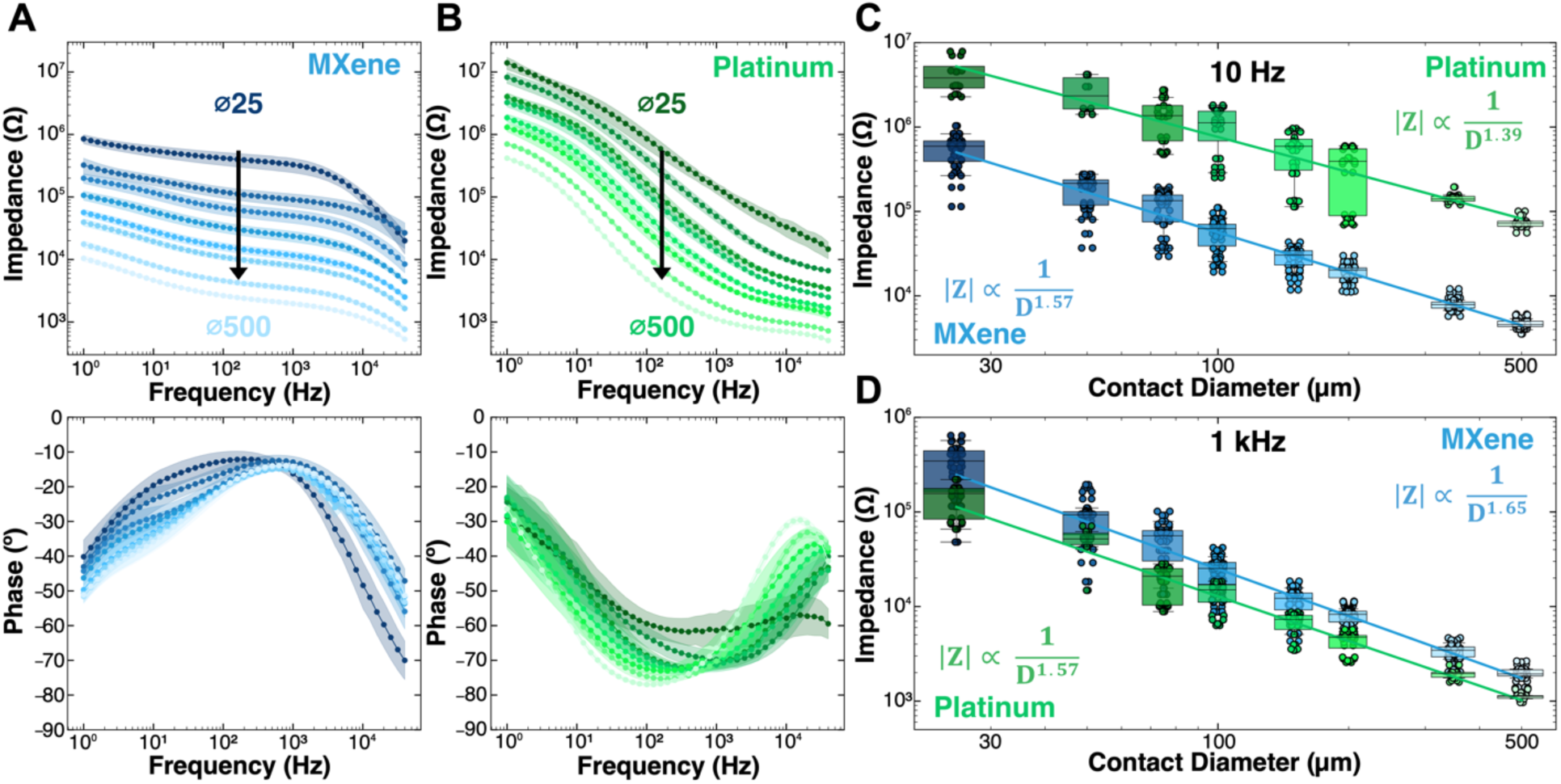
Electrochemical impedance spectroscopy of Ti_3_C_2_T*_x_* MXene and platinum microelectrodes at varying diameters. **(A)** Impedance modulus and phase spectra of Ti_3_C_2_T_x_ (n_MX_ = 34 contacts per diameter) and **(B)** Pt microelectrodes of varying diameter (n_Pt_ = 18 contacts per diameter). Individual points represent mean, shaded areas represent ± 1 standard deviation. **(C, D)** Impedance modulus at **(C)** 10 Hz (R^2^_MX, 10 Hz_ = 0.93, R^2^_Pt, 10 Hz_ = 0.78) and **(D)** 1 kHz (R^2^_MX, 1 kHz_ = 0.92, R^2^_Pt, 1 kHz_ = 0.93) as a function of electrode diameter.

For all measured diameters, we found that at 10 Hz the impedance modulus of Ti_3_C_2_T*_x_* microelectrodes was roughly an order of magnitude lower than that of Pt microelectrodes (Ti_3_C_2_T*_x_*: 500 µm |**Z_10 Hz_**| = 4.67 ± 0.58 kΩ; 25 µm: |**Z_10 Hz_**| = 550 ± 218 kΩ; Pt: 500 µm |**Z_10 Hz_**| = 74.40 ± 12.20 kΩ; 25 µm: |**Z_10 Hz_**| = 4.37 ± 1.96 MΩ; all other values in **Supplementary Table S1**). At 1 kHz, the impedance modulus for the two materials is almost equivalent (Ti_3_C_2_T*_x_*: 500 µm |**Z_1 kHz_**| = 1.97 ± 0.34 kΩ; 25 µm: |**Z_1 kHz_**| = 328 ± 168 kΩ; Pt: 500 µm |**Z_1k_**| = 1.11 ± 0.11 kΩ; 25 µm: |**Z_1 kHz_**| = 140 ± 74.70 kΩ; all other values in **Supplementary Table S1**). The size-dependency of the impedance modulus at 10 Hz and 1 kHz follows a power-law relationship (|**Z**| ∝ 1/D*^x^*) where D is contact diameter and *x* is an exponent that varies between 0 and 2, with *x* = 1 representing a perimeter dependency and *x* = 2 representing an area dependency (**Figure 2C, 2D**).^64^ For both Ti_3_C_2_T*_x_* and Pt, the fitted impedance modulus as a function of the electrode diameter at 1 kHz shows a sub-areal dependency: 1/D^1.65^ for Ti_3_C_2_T*_x_* and 1/D^1.57^ for Pt, consistent with prior reports for Pt microelectrodes (1/D^1.64^, 1/D^1.70^).^64^ This was consistent in the fitted impedance moduli at 10 Hz as well for both materials. These intermediate exponents are indicative of a combination of both planar and spherical diffusion of ions occurring at the electrode-electrolyte interface, where planar diffusion dominates in large electrodes and spherical diffusion dominates as the electrode approximates and then becomes smaller than the diffusion layer.^64^ Ultimately, from this analysis, the fitted impedance slopes at 1 kHz and 10 Hz collected for each material can be used to estimate the impedance magnitude and potential SNR tradeoff for future microelectrodes across a range of electrode diameters.

### Impedance modeling informs charge transfer properties

To better understand the mechanisms governing the electrochemical charge transfer occurring between the thin-film microelectrodes and the ionic medium, we first qualitatively inspected the Nyquist plots of the two materials before turning to quantitative fitting with equivalent circuit models (**Figure 3A, S3**). Notably, at sufficiently high frequencies (> 300 Hz, denoted by a distinct semicircle for both materials), charge transfer responses are known to be dominated by capacitive and kinetically-controlled reactions; and for Ti_3_C_2_T*_x_* MXene microelectrodes, this corresponds primarily to capacitive double layer charging and discharging, with some contributions of faradaic processes involving hydrogen-based surface terminations (e.g., -OH^-^, H^+^).^65–67^ These dynamics are typically quantified using a single electrochemical time constant *τ* = RC, where R is the charge-transfer resistance and C is the double-layer capacitance. For Ti_3_C_2_T*_x_* microelectrodes, however, the complex Nyquist profile suggests the presence of two distinct time constants, each corresponding to separate RC pairs in the equivalent circuit, reflected as isolated semicircles in the Nyquist plot.^67^ The first time constant, *τ*_1_, observed at higher frequencies, reflects rapid surface-level capacitive double layer formation and electron transfer kinetics. As the frequency decreases, however, slower ion diffusion processes begin to dominate, leading to an abrupt ∼45° angle in the Nyquist plot. Concurrent to this diffusion mechanism is the emergence of a second, broader semicircle in the Nyquist plot around 10 Hz (τ_2_), representing a significantly larger time constant (*τ*_2_ >> *τ*_1_). This second RC response indicates deeper double layer charging/discharging and kinetic charge transfer behavior within the layered architecture of the Ti_3_C_2_T*_x_* film. At very low frequencies (< 2 Hz), finally, there is a steep uptrend in the slope of the Nyquist plot reflecting that diffusion-dominated ionic transport now governs the electrochemical response throughout the entire thickness of the Ti_3_C_2_T*_x_* film, both at the surface and through the overlapping flakes. Notably, this multi-timescale behavior is unique to Ti_3_C_2_T*_x_* microelectrodes, as conventional Pt electrodes – composed of continuous, crystalline films which are impermeable to ions – exhibit only surface-confined electrochemical activity and a single dominant time constant (**Figure 3B, S4**).

**Figure 3.**
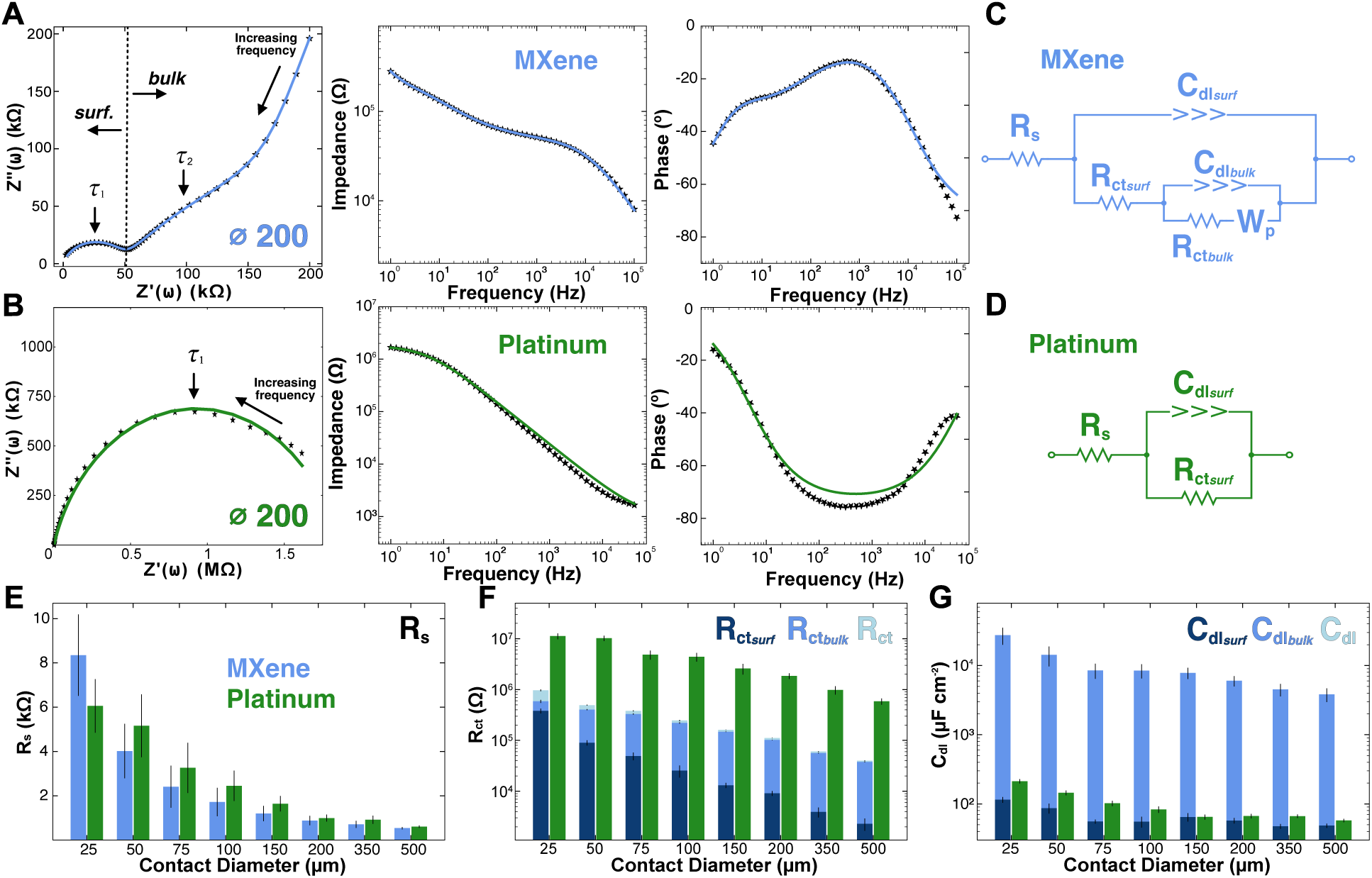
Ti_3_C_2_T*_x_* and Pt equivalent circuit modeling. (A, **B)** Nyquist, impedance, and phase plots of 200 µm **(A)** Ti_3_C_2_T*_x_* and **(B)** Pt microelectrodes. Markers are data points, and continuous curves are the fitted models. **(C, D)** Equivalent circuit models for **(C)** Ti_3_C_2_T*_x_* **(D)** Pt microelectrodes. **(E)** Comparative series resistance (R*_s_*). **(F)** Charge transfer resistance of Ti_3_C_2_T*_x_* (blue) and Pt (green). Ti_3_C_2_T*_x_*: R*_ct_* = R*_ct, surf_* + R*_ct, bulk_*; Pt: R*_ct_* = R*_ct, surf_* . **(G)** Double-layer capacitance of Ti_3_C_2_T*_x_* (blue) and Pt (green). Ti_3_C_2_T*_x_*: C*_dl_* = C*_dl, surf_* + C*_dl, bulk_*; Pt: C*_dl_* = C*_dl, surf_*. Note that the cumulative C*_dl_* is dominated by the bulk capacitance, and it is not visible at the scale of the plot.

By using the qualitative analysis of the multiple time- and frequency-dependent responses of the charge transfer processes of the Ti_3_C_2_T*_x_* microelectrodes in addition to our understanding of the layered nature of the spray-coated Ti_3_C_2_T*_x_* films, we next fitted the EIS data for both materials with physically-informed equivalent circuit models. Each element of the circuit represents a unique process taking place at the electrode-electrolyte interface, which can be categorized into solution interactions, kinetic and double layer charging and discharging, and diffusion interactions (**Figure 3C**). Briefly, the solution interactions are represented by the solution resistance (R*_s_*), which is primarily controlled by the electrolyte conductivity (𝜌) and – for circular electrodes – by the electrode radius (**r**, **Equation 1**).^68^

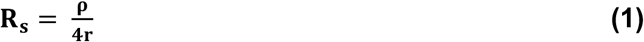

The double layer and kinetic charge transfer interactions at the surface layers of the electrode are then represented by a simplified Randles circuit, consisting of the interfacial (or surface) charge-transfer resistance (R*_ct, surf_*) in parallel with the interfacial double-layer capacitance (C*_dl, surf_*). For the Ti_3_C_2_T*_x_* microelectrodes, however, there is an additional RC nested subcircuit representing the double-layer (C*_dl, bulk_*) and kinetic charge-transfer reactions (R*_ct, bulk_*) occurring within the bulk of the layered architecture of the film, as well as a semi-infinite porous Warburg element (W*_p_*). These nested elements of the Ti_3_C_2_T*_x_* microelectrode’s equivalent circuit model account for the slower intercalation of ions penetrating the layered film architecture at low frequencies (<100 Hz).^69^ The electrochemical interactions occurring at the electrode/electrolyte interface of Pt microelectrodes can be modeled by a single Randles circuit without any intercalation or diffusion interaction. (**Figure 3D**). In both models, all capacitive elements are represented as constant phase elements (CPEs), which better reflect real-world capacitive behavior by accounting for surface roughness and non-ideal electrode properties, including impurities.^70^ CPE parameters are converted into conventional capacitance values using **Equation 2**, where Y_o_ and α are the characteristic parameters of the CPE, and ω is the angular frequency at which the imaginary component of the impedance reaches its maximum in the Nyquist plot.

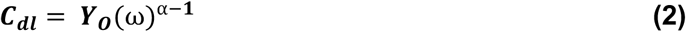

To first validate that the measured EIS spectra satisfied linearity and causality assumptions and to ensure the models sufficiently parameterize the charge-transfer processes,^71^ we performed Kramers-Kronig (KK) consistency checks (Lin–KK) and frequency-resolved residual/error analysis of the equivalent circuit fit, confirming KK consistency from 1 Hz to 10 kHz (**Figure S5**). Comparing the values obtained from equivalent circuit modeling (i.e., R*_s_*, R*_ct_*, C*_dl_*) reveals significant differences in the charge transfer mechanisms between Ti_3_C_2_T*_x_* and Pt microelectrodes across all sizes (**Table S2, S3**). The solution resistance (R*_s_*) is nearly identical between the two materials for all the electrode diameters: 500 µm, **R_s, MX_** = 540 ± 53 Ω, **R_s, Pt_** = 603 ± 55 Ω; 25 µm: **R_s, MX_** = 8.15 ± 1.84 kΩ, **R_s, Pt_** = 7.06 ± 1.21 kΩ, (**Figure 3E**). The cumulative charge transfer resistance (Ti_3_C_2_T*_x_*: R*_ct_ = R_ct, surf_ + R_ct, bulk_*; Pt: R*_ct_ = R_ct, surf_*) of the two materials, however, differs drastically: Ti_3_C_2_T*_x_* electrodes exhibit over an order of magnitude lower charge-transfer resistance compared to platinum (**Figure 3F**). Furthermore, as the microelectrode diameter decreases, as expected, the charge-transfer resistance increases, due to the reduced surface area available for ionic exchange.^72^ This is directly grounded in the formula for charge-transfer resistance shown in **Equation 3**, where k^0^ is the electron transfer rate, n is the number of electrons transferred in the reaction, F is Faraday’s constant, R is the gas constant, T is the temperature, C is the concentration of the redox species, and A is the surface area of the electrode.

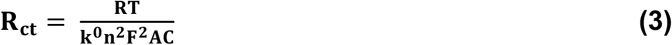

Empirically, the cumulative charge-transfer resistances for the Ti_3_C_2_T*_x_* and Pt microelectrodes are: 500 µm, **R_ct, MX_** = 39.90 ± 6.15 kΩ, **R_ct, Pt_** = 585 ± 83.30 kΩ; 25 µm, **R_ct, MX_** = 966 ± 209 kΩ, **R_ct,Pt_** = 11.20 ± 1.46 MΩ. Similarly, the cumulative double-layer capacitance (C*_dl_ = C_dl, surf_ + C_dl, bulk_*) of Ti_3_C_2_T*_x_* microelectrodes is roughly two orders of magnitude higher than that of Pt for all contact sizes (**Figure 3G**): 500 µm, **C_dl, MX_** = 3,820 ± 851 µF cm^-2^, **C_dl, Pt_** = 57.50 ± 3.23 µF cm^-2^; 25 µm, **C_dl, MX_** = 27.60 ± 7.59 mF cm^-2^, **C_dl, Pt_** = 0.21 ± 0.01 mF cm^-2^. This enhanced capacitive behavior in the MXene compared to platinum microelectrodes arises from two key factors: (1) the layered flake architecture of the Ti_3_C_2_T*_x_* films, which permits electrolyte penetration into the bulk of the contacts and increases accessible surface area,^73^ and (2) the high intrinsic capacitive behavior of Ti_3_C_2_T*_x_*, which reflects not only electrostatic double-layer charging but also contributions from fast, reversible redox (pseudocapacitive) processes associated with surface terminations and ion intercalation.^74^ These mixed mechanisms further explain why the phase response of the MXene electrodes is not purely capacitive, but instead shows faradaic contributions, consistent with pseudocapacitive charge storage observed in prior studies.^65–67^

### Cyclic voltammetry and charge storage capacity analysis

Cyclic voltammetry (CV) is used to analyze surface reactions at the electrode-electrolyte interface, assess reaction reversibility, and quantify electrochemical capacitance, where the current response in a voltammogram reflects both capacitive double layer charging and faradaic charge transfer processes.^75,76^ CVs were collected in 1x PBS at 500 mV s^-1^ vs a Ag/AgCl reference on each contact size for Ti_3_C_2_T*_x_* and Pt devices. The potential window for the thin-film Ti_3_C_2_T_x_ microelectrodes is -1.55 V to 0.35 V vs Ag/AgCl in PBS, which was experimentally determined (**Figure S6**) and closely matches previously established voltage limits for Ti_3_C_2_T_x_.^55^ The Pt potential window is well-known and ranges from -0.60 V to 0.80 V vs Ag/AgCl in PBS. As expected, for both Ti_3_C_2_T*_x_* and Pt microelectrodes, the measured current decreased with electrode size (**Figure 4A, B**). Specifically, the cathodic charge stored for Ti_3_C_2_T*_x_* microelectrodes was: 500 µm, **Q_st_** = 40.80 ± 11.40 µC; 25 µm, **Q_st_** = 0.59 ± 0.48 µC. For Pt: 500 µm, **Q_st_** = 1.31 µC ± 0.11 µC; 25 µm, **Q_st_** = 0.02 µC ± 0.01 µC (**Figure 4C**). As such, Ti_3_C_2_T*_x_* microelectrodes show ∼50x more charge stored compared to size-matched Pt across all diameters. This, in addition to the largely rectangular shape of the Ti_3_C_2_T*_x_* voltammograms and in agreement with the impedance modeling, supports the highly capacitive nature of the Ti_3_C_2_T*_x_* flakes assembled in the microelectrode form. Although the voltammogram of the Ti_3_C_2_T*_x_* microelectrodes is dominated by double layer capacitive charging, two distinct faradaic features are observed near 0 V and -0.83 V vs. Ag/AgCl. These peaks are attributed to hydrogen-based redox processes associated with surface terminations – specifically, proton reduction/oxidation and hydroxyl-related reactions consistent with the H⁺ and -OH^-^ groups present on Ti_3_C_2_T*_x_* surfaces.^74,77,78^ As contact diameter decreases, however, both charge-transfer resistance and the area-normalized double-layer capacitance increase, causing the faradaic peaks to shrink, leading to a nearly perfectly capacitive voltammogram for the smallest 25 µm contacts. This can be physically understood as the charge transfer mechanisms shifting from planar, surface-dominating faradaic reactions at larger contact sizes to more radial, layered, bulk-based capacitive reactions in the smaller contacts as the volume to surface area ratio increases with decreasing contact diameter.^79,80^

**Figure 4.**
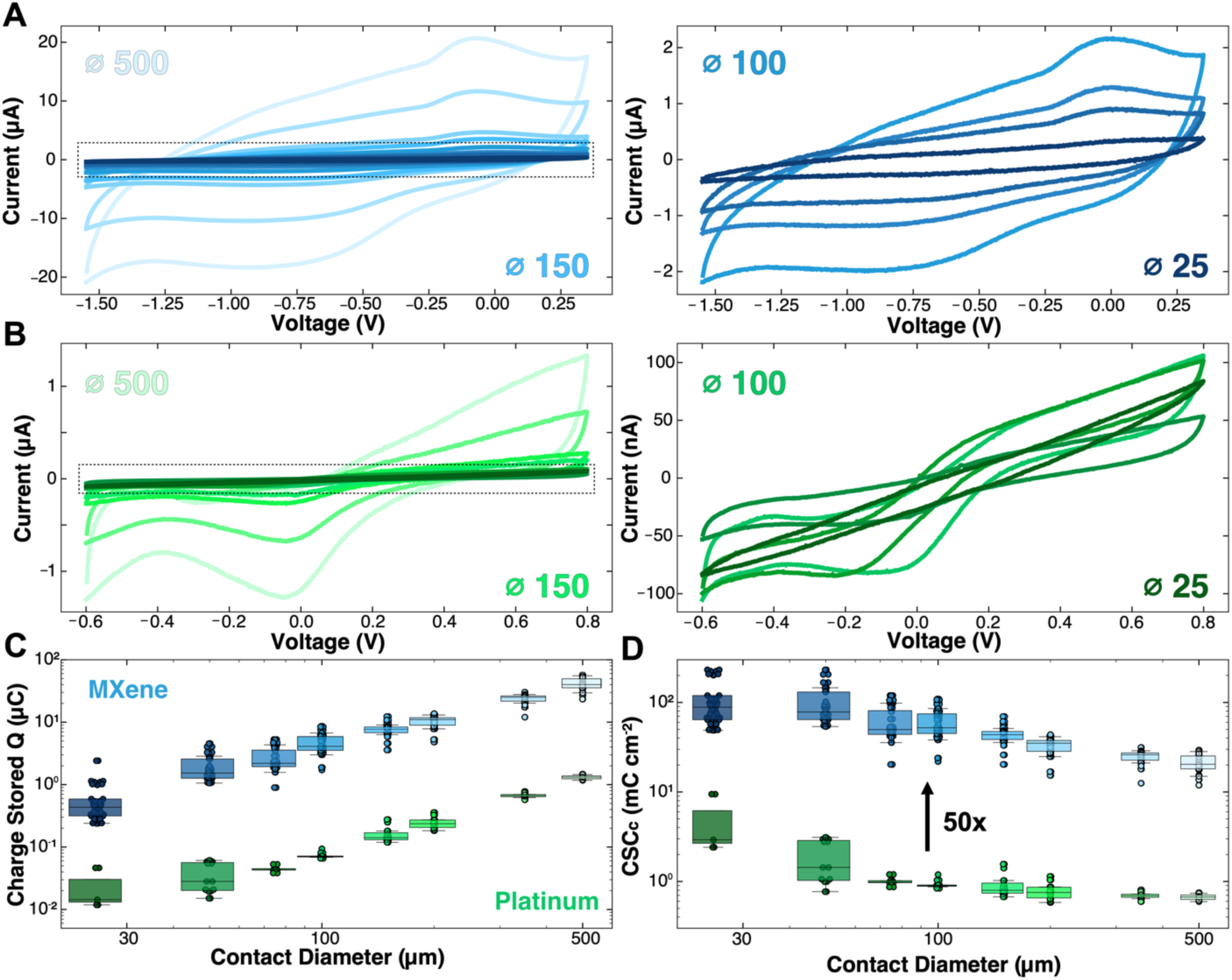
Cyclic voltammetry and charge storage capacity analysis. **(A)** *Left:* Averaged cyclic voltammograms of Ti_3_C_2_T*_x_* MXene microelectrodes from -1.55 V (cathodic limit) to 0.35 V(anodic limit). *Right:* Zoomed-in voltammograms for contacts ranging from 100 µm to 25 µm (n_MX_ = 34 contacts per diameter). **(B)** *Left:* Averaged cyclic voltammograms of sputtered Pt microelectrodes from -0.6 V (cathodic limit) to 0.8 V(anodic limit). *Right:* Zoomed-in voltammograms for contacts ranging from 100 µm to 25 µm (n_Pt_ = 18 contacts per diameter). **(C)** Charge stored across all contact sizes for Ti_3_C_2_T*_x_* and Pt devices. **(D)** Cathodic charge storage capacity across all contact sizes of Ti_3_C_2_T*_x_* and Pt devices.

In stimulation electrodes, the measured capacitance is particularly important as it governs the material’s ability to store and deliver charge efficiently. Specifically, the cathodal charge storage capacity (CSCc) provides a measure of how much charge can be safely stored at the surface and throughout the bulk of the electrode contact, with higher CSCc values corresponding to more effective stimulation performance.^12,13^ The CSCc of the microelectrodes was calculated for each contact size by normalizing the cathodal charge calculated by the contact’s surface area. From this calculation, we found that the smallest microelectrode contacts had the largest CSCc for both Ti_3_C_2_T*_x_* and Pt devices, with Ti_3_C_2_T*_x_* microelectrodes showing ∼50x higher CSCs than the Pt microelectrodes for all sizes: Ti_3_C_2_T*_x_* 500 µm, **CSCc** = 20.80 ± 5.83 mC cm^-2^; 25 µm, **CSCc** = 119.90 ± 97.30 mC cm^-2^; Pt 500 µm: **CSCc** = 0.67 ± 0.05 mC cm^-2^; 25 µm: **CSCc** = 4.29 ± 2.99 mC cm^-2^ (**Figure 4D, Table S4**). This behavior is caused by the increase in capacitive double layer reactions compared to faradaic reactions as the thickness-to-surface area ratio increases with decreasing electrode diameter.

CVs can also provide a quantification of the material’s total double layer capacitance, referred to as the ‘specific capacitance’. This measurement is conducted by acquiring CV in a limited voltage range (here, between -0.3 V and 0.3 V) at increasing scan rates from 50 mV s^-1^ to 500 mV s^-1^, and then extracting the half-cell current at 0V for each independent scan rate (( I_0v,A_ – I_0v,C_ ) / 2) (**Figure S7A**). The slope of the linear fit of the current Vs. the scan rate is the specific capacitance (**Figure S7B**). From this measurement, we found that the area-normalized specific capacitance increases with decreasing contact diameter: Ti_3_C_2_T*_x_* 500 µm, 4,060 ± 942 µF/cm^2^; 25 µm, 19,300 ± 7,640 µF/cm^2^ (**Figure S7C**); Pt 500 µm, 49.20 ± 3.85 µF/cm^2^; 25 µm, 249 ± 21.10 µF/cm^2^.

Importantly, the measured values of the specific capacitance are in agreement with the fitted C_dl_ in the equivalent impedance model (**Table S5**), confirming the significantly larger contribution of capacitive mechanisms of charge transfer at the electrode-electrolyte interface for Ti_3_C_2_T*_x_*, especially for electrode diameters < 100 µm.

### Voltage transients and charge injection capacity analysis

Voltage transient analysis is a fundamental technique for assessing the performance of implantable electrodes for electrical stimulation. By monitoring the voltage response to controlled current pulses, key parameters such as electrode polarization behavior, charge injection limits, electrochemical stability, and implantable pulse generator (IPG) requirements can be determined. The maximum cathodic (E_mc_) and anodic (E_ma_) polarization potentials define whether the electrode operates within the safe potential window, beyond which adverse reactions such as water electrolysis and gas evolution may occur. Identifying the onset of these limits enables calculation of the maximum safe current amplitude and the charge injection capacity, a critical metric quantifying the maximum safely injectable charge per phase.^13,81^

In the current study, we pulsed the Ti_3_C_2_T*_x_* and Pt microelectrodes at increasing current amplitudes until the maximum cathodic polarization potential (E_mc_) reached the cathodic potential limit for each contact size (**Figure 5A**). The anodic polarization potential never reached the anodic limit first in any of our tests. For all voltage transients described henceforth for both MXene and platinum microelectrodes, the bias voltage (E_ipp_) was recorded Vs. an Ag/AgCl reference, the cathodic pulse-width was held at 500 µs, and the anodic pulse-width was held fixed at 1000 µs at half the amplitude of the cathodic pulse to maintain a charge-balanced waveform. The maximum cathodic current limits for Ti_3_C_2_T*_x_* microelectrodes decreased with the contact diameter, 500 µm, **I_max_** = 1884.10 ± 200.10 µA; 25 µm, **I_max_** = 4.88 ± 1.90 µA. We observed a similar trend for Pt microelectrodes, although the maximum current limits were significantly smaller: 500 µm, **I_max_** = 133.70 ± 8.35 µA; 25 µm, **I_max_** = 1.38 ± 0.55 µA (**Figure 5B**).

**Figure 5.**
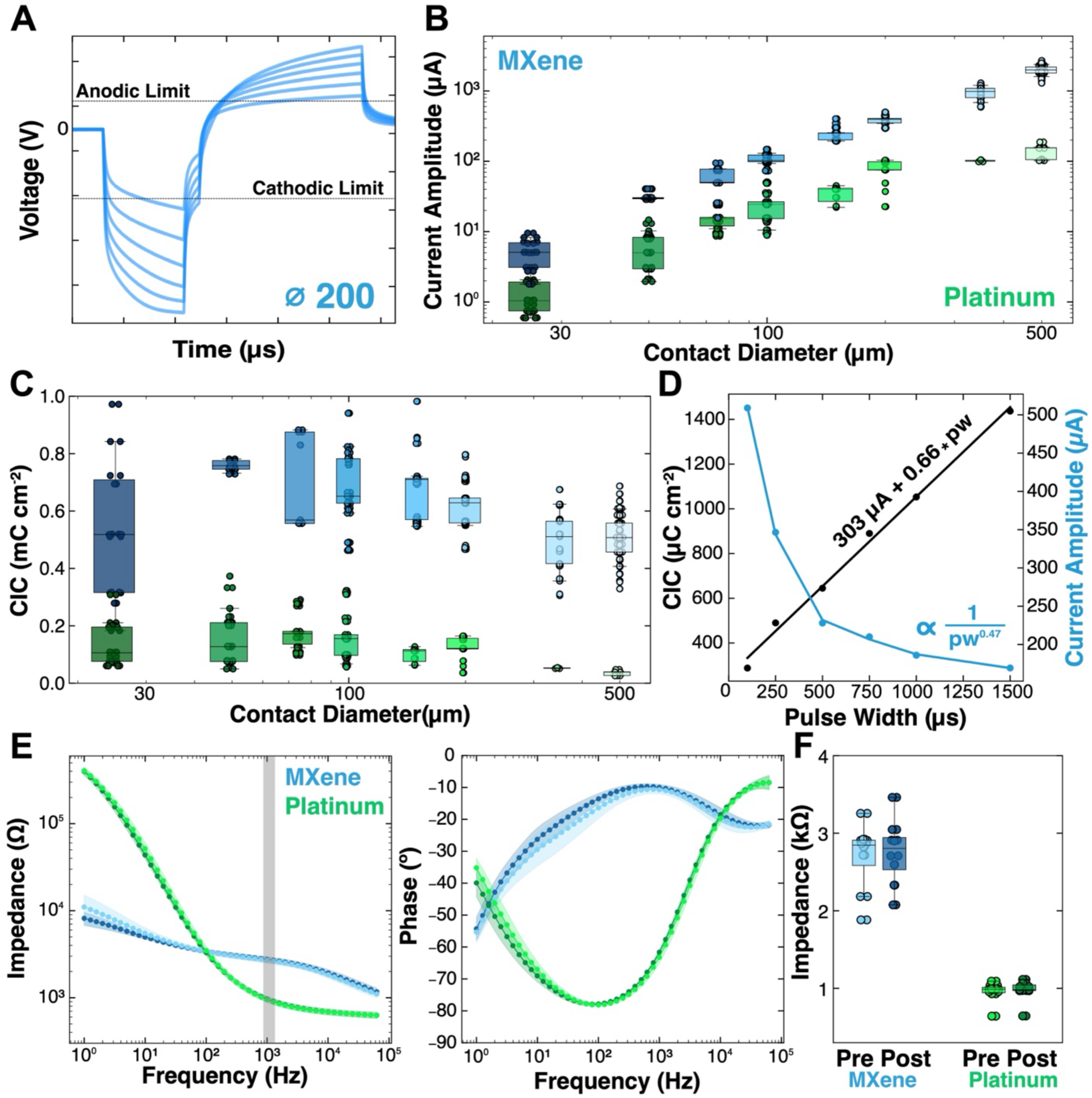
Voltage transients, charge injection capacity, effects of pulse width, and continuous stimulation. **(A)** Representative voltage transients at increasing current injections approaching the cathodic potential window limit for 200 µm Ti_3_C_2_T*_x_* microelectrodes. **(B)** Maximum cathodic current limit for Ti_3_C_2_T*_x_* and Pt microelectrodes for contact sizes. (n_MX_ = 34 contacts per diameter, n_Pt_ = 18 contacts per diameter). (**C**) Cathodic charge injection capacity for all Ti_3_C_2_T*_x_* and Pt microelectrodes. (**D**) Fitted cathodic charge injection capacity and maximum current limit as a function of cathodic pulse-widths for a 150 µm Ti_3_C_2_T*_x_* microelectrode. CICc = 0.80 pw + 253 µC cm^-2^, R^2^ = 0.99; I_max_ ∝ 1 / pw^0.42^, R^2^ = 0.98. (**E**) EIS magnitude and phase for 500 µm Ti_3_C_2_T*_x_* and Pt microelectrodes before (light) and after (dark) 5000 continuous stimulation pulses at 100 Hz (n = 8 electrodes, each). (**F**) 1 kHz impedance modulus pre (light) and post (dark) stimulation.

The cathodic charge injection capacity (CICc) for both Ti_3_C_2_T*_x_* and platinum microelectrodes remained relatively stable with the electrode diameter, with the smallest microelectrodes showing slightly larger CICc values. This trend of CICc vs surface area is consistent with what others have noted in literature;^34,82^ however, the lack of pronounced edge effects noted at different contact sizes in voltage transients is not necessarily indicative of how the stimulation interacts with the body during *in-vivo* microstimulation, where edge profiles can greatly affect the distance and intensity of stimulation.^80^ Ultimately, the averaged CICc for Ti_3_C_2_T*_x_* microelectrodes is consistently ∼6x greater than the CICc of Pt microelectrodes across all sizes (CICc: 592.40 ± 112.40 µC/cm^2^, Pt: 111.40 ± 33.30 µC/cm^2^) (**Figure 5C**). Additionally, it is worth noting that the CICc for the sputtered Pt is slightly lower than in other reports due to the lack of a positive bias voltage, which was not conducted in this study to ensure an even comparison with the Ti_3_C_2_T*_x_* microelectrodes (**Table S6**).^34^

Further current limit and CICc characterization were conducted at several other pulse widths, with the cathodic pulse-width ranging from 100 µs to 1500 µs for all contact sizes for both Ti_3_C_2_T*_x_* and Pt devices (**Table S7, S8**). By increasing or decreasing the pulse-width, the maximum injectable current decreases or increases, respectively. The change in maximum current amplitude is not linear, but rather decays exponentially with the pulse-width (**I_max_** ∝ 1/pw^x^ ) where pw is the cathodic pulse-width and x is a non-negative value. CICc, however, increases linearly with pulse-width (**CICc** = a*pw + b) where a and b are constants (**Figure 5D**). For a 150 µm Ti_3_C_2_T*_x_* electrode contact, **I_max_** ∝ 1/pw^0.47^, R^2^ = 0.98 and **CICc** = 0.66*pw + 303 µC cm^-2^, R^2^ = 0.99. These results, repeated with a near-identical trend in a similar study of PEDOT: PSS / Au microelectrodes,^34^ allow for a precise understanding of the inverse tradeoff of the Ti_3_C_2_T*_x_* microelectrodes’ maximum stimulation current amplitudes Vs. injectable CIC at any given pulse-width – critical for optimizing application-specific stimulation waveforms. Finally, each 500 µm diameter Ti_3_C_2_T*_x_* and Pt microelectrode was subjected to 5000 pulses of 250 µs, biphasic, charge-balanced, 100 Hz stimulation trains at a sub-CIC threshold amplitude of 350 µA. Before and after impedance measurements showed the lack of a significant change in the impedance modulus at 1 kHz caused by the stimulation (*p_MX_* = 0.78, *p_Pt_* = 0.79, **Figure 5E, F**).

### Effects of Ti_3_C_2_T*_x_* concentration on the electrochemical behavior of the microelectrodes

Briefly, three separate groups of 200 µm Ti_3_C_2_T*_x_* microelectrodes were identically fabricated while varying only the concentration of the solution used for the film fabrication: 1 mg mL^-1^, 3 mg mL^-1^, or 5 mg mL^-1^ Ti_3_C_2_T*_x_* MXene. The spray-coated volume was kept constant at 50 mL. AFM showed that increasing the concentration of Ti_3_C_2_T*_x_* results in an increase in surface roughness, which was also observed for spray-cast electrodes (1 mg mL^-1^: roughness_avg_ = 10.3 nm, range = 146 nm, 3 mg mL^-1^: roughness_avg_ = 16.9 nm, range = 213 nm, 5 mg mL^-1^: roughness_avg_ = 21.1 nm, range = 271 nm, n = 4 devices (8 contacts), each condition, **Figure 6A**). Although a higher surface roughness did not appear to greatly affect the overall magnitude of impedance (**Figure 6B**), its effect was evident in the fitted equivalent circuit parameters (**Figure 6C**). Specifically, rougher electrodes had the lowest R_ct_, likely due to the increased accessible surface area available for faradaic processes at the charged surface groups. Similarly, C_dl_ and CSCc measurements (**Figure 6D**) show larger capacitance as the roughness of the microelectrode contact increases, as there are more flake layers and locations with higher charge density on the smoother surfaces (**Figure 6E**). The increased film thickness resulting from the higher Ti_3_C_2_T*_x_* concentration was confirmed with profilometry of the three conditions (1 mg mL^-1^: 265 ± 53 nm, 3 mg mL^-1^: 932 ± 208 nm, 5 mg mL^-1^: 1,895 ± 152 nm, **n** = 15 measurements). The voltage transient and current excursions, and the calculated CICc, did not appear to be affected by the Ti_3_C_2_T*_x_* concentration (**Figure 6F, G**). Ultimately, from the collected measurements, it is clear that both impedance measurements to quantify recording capabilities and voltage transients to quantify stimulation capabilities are largely area dependent and not significantly affected by a change in surface roughness or film thickness within the tested range of spray-coated concentration of Ti_3_C_2_T*_x_* MXene.

**Figure 6.**
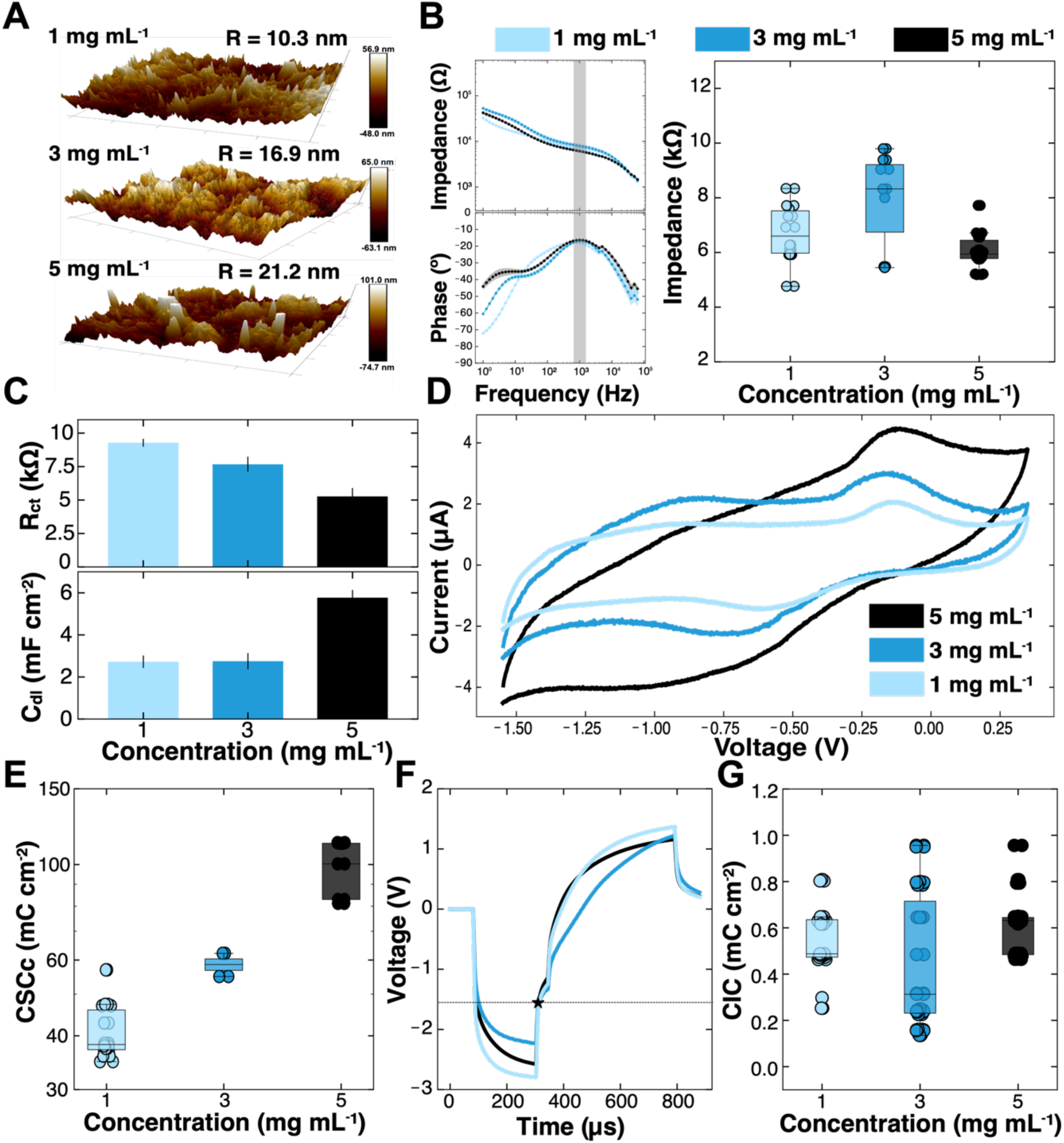
Effects of the concentration of the Ti_3_C_2_T*_x_* solution. **(A)** AFM surface profiles for *Top:* 1 mg mL^-1^, *Middle:* 3 mg mL^-1^, *Bottom:* 5 mg mL^-1^ Ti_3_C_2_T*_x_* spray-coated microelectrodes. (**B**) *Left:* impedance and phase spectra. *Right:* 1 kHz impedance magnitude for 200 µm microelectrode fabricated with 50 mL of 1, 3, and 5 mg mL^-1^ of Ti_3_C_2_T*_x_*. **(C)** Circuit-fitted model magnitudes for charge transfer resistance and double layer capacitance as a function of spray-coated concentration. **(D)** CV of each spray-coated concentration. **(E)** Charge storage capacity as a function of sprayed concentration for 200 µm MXene contact. **(F)** Averaged maximum voltage transient for each spray-coated concentration. **(G)** Maximum cathodic charge injection capacity as a function of sprayed concentration. N = 8 electrode contacts for all measurements.

### Effects of the spray-coated volume on the electrochemical behavior of Ti_3_C_2_T*_x_* microelectrodes

For this study, all 200 µm microelectrode contacts were fabricated using the 3 mg mL^-1^ Ti_3_C_2_T*_x_* solutions at the following volumes: 25 mL, 50 mL, 75 mL, and 100 mL. As expected, the main effect of varying the spray-coated volume is in the film thickness: 25 mL, 209.70 ± 8.80 nm; 50 mL: 478 ± 100.90 nm; 75 mL: 804 ± 18 nm; 100 mL: 1067 ± 93.20 nm, n = 4 devices (8 contacts), each (**Figure 7A**). Similar to what we observed with the variation in Ti_3_C_2_T*_x_* concentration, the main effect of increasing the solution volume was the increase in double-layer capacitance, likely caused by an increased access to capacitive charge-discharge reactions throughout the thicker Ti_3_C_2_T*_x_* film (**Figures 7B, C**). In line with the increased thickness and capacitance, it was clear in the CVs and the calculated CSCc that as the film thickness increased with increasing volume, so too did the width of the voltammogram and the magnitude of the CSCc (**Figure 7D, E**). Once again, however, similar to what was shown as a result of varying the sprayed concentration, there was a nearly non-existent change in the voltage transients or the calculated CICc as a function of varying the volume sprayed for the MXene microelectrodes (**Figure 7F, G**). Here again we see that although the mechanisms of interaction between the device and the interface were slightly altered by the variation in sprayed volume, predominantly in the C_dl_ and CSCc, there was little to no change in the recording or stimulation capabilities of the device by increasing or decreasing the volume, further confirming that recording and stimulation functionality is overwhelmingly controlled by the geometric surface area.

**Figure 7.**
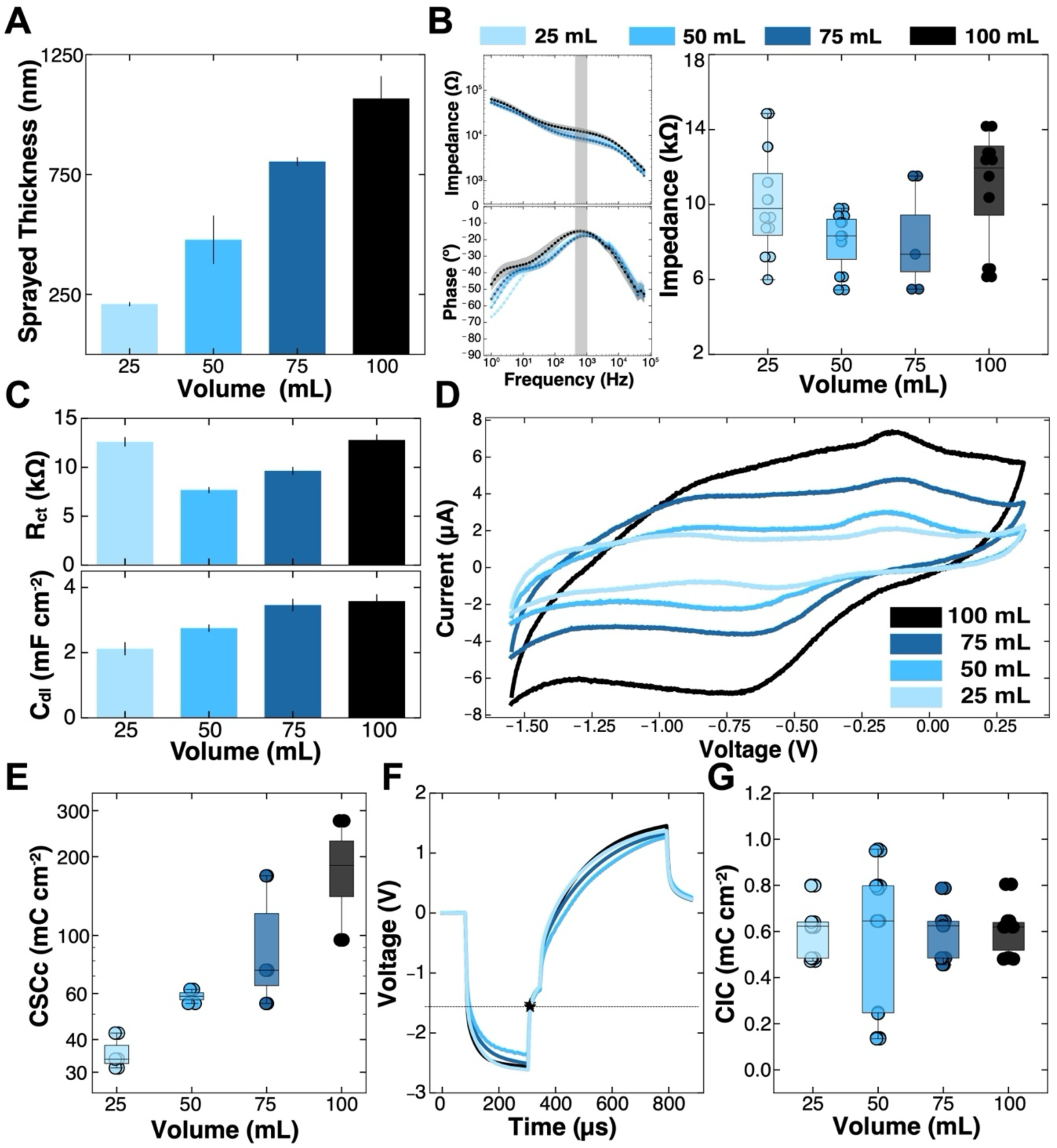
Effects of Ti_3_C_2_T*_x_* volume. **(A)** Film thickness for Ti_3_C_2_T*_x_* microelectrode fabricated from 3 mg mL^-1^ solutions at varying volumes. (**B**) *Left:* impedance and phase spectra. *Right:* 1 kHz impedance magnitude of 200 µm microelectrodes c. **(C)** Fitted R_ct_ and C_dl_, **(D)** cyclic voltammograms, and **(E)** charge storage capacity at varying Ti_3_C_2_T*_x_* volumes. **(F)** Averaged maximum voltage transient for each spray-coated volume. **(G)** Maximum cathodic charge injection capacity. N = 8 electrode contacts for all measurements.

## DISCUSSION

The electrochemical response of Ti_3_C_2_T*_x_* microelectrodes across microscale dimensions highlights how a combination of intrinsic material properties and accessible surface area jointly determine the device performance for electrical recording and stimulation. Through impedance modeling, cyclic voltammetry, and voltage transient analysis, this work reveals a consistent mechanistic picture: the Ti_3_C_2_T*_x_* films’ layered morphology and abundant surface terminations promote rapid ion exchange and mixed capacitive and pseudocapacitive charge storage, resulting in low interfacial resistance and high effective capacitance across diameters ranging from 25 µm - 500 µm.^73^ These findings advance the work of previous studies on MXene-based bioelectronics by quantitatively resolving how charge-transfer pathways evolve with electrode miniaturization and film morphology - an essential step toward rationally engineering electrode-electrolyte interfaces for high-density neural stimulation and recording.^39,55,60^

By analyzing key parameters in the impedance model of Ti_3_C_2_T*_x_* microelectrodes, it becomes clear that the electrochemical exchange with the electrolyte is governed mostly by capacitive processes captured by large C_dl_ with kinetic contributions reflected in R_ct_. Comparing equivalent-circuit values (R_s_, R_ct_, C_dl_) revealed that R_s_ is similar for MXene and Pt at all given diameters, as expected for identical geometries and electrolytes, whereas the cumulative R_ct_ is over ten times lower and the cumulative C_dl_ is roughly two orders of magnitude higher for Ti_3_C_2_T*_x_* across all diameters. This behavior is likely caused by both the layered film architecture of the Ti_3_C_2_T*_x_* microelectrodes enabling electrolyte access beyond the immediate surface, as well as from the high intrinsic capacitance of the Ti_3_C_2_T*_x_* flakes (**Figure 8**). These circuit-level differences explain the stable, low-magnitude impedance at lower frequencies (<1000 Hz) in the EIS measurements and the broader, more capacitive CVs observed for MXene compared with Pt.

**Figure 8.**
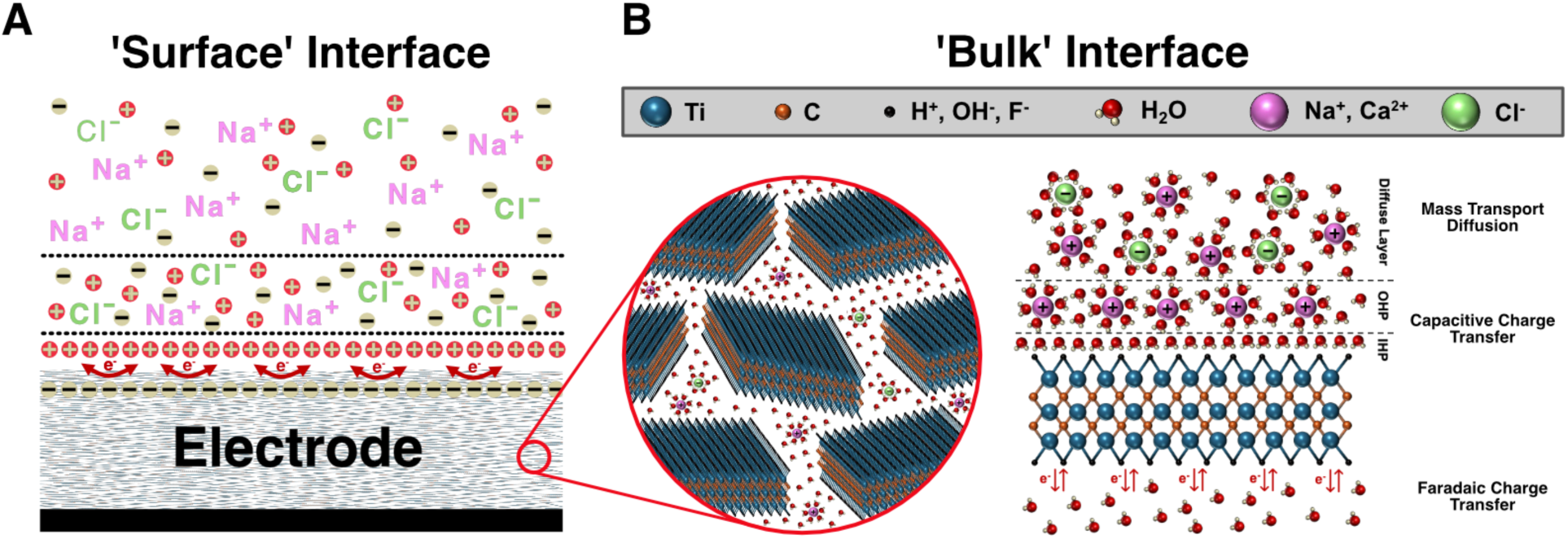
Schematic diagram of the charge-transfer mechanisms of Ti_3_C_2_T_x_ MXene. **(A)** Faradaic and capacitive charge-transfer at the surface interface of the Ti_3_C_2_T_x_ MXene microelectrodes. **(B)** Zoomed-in view of the faradaic, double-layer capacitive, and mass-transport diffusion processes occurring within the layered bulk of the film. Faradaic redox reactions stemming from T_x_ groups of Ti_3_C_2_T_x_ flakes (OH^-^, H^+^, F^-^). Capacitive charge-transfer from Inner-and Outer Helmholtz Planes (IHP, OHP). Diffusion comes from the mass transport of larger ionic species. Figure 8**. Schematic diagram of the charge-transfer mechanisms of Ti_3_C_2_T_x_ MXene. (A)** Faradaic and capacitive charge-transfer at the surface interface of the Ti_3_C_2_T_x_ MXene microelectrodes. **(B)** Zoomed-in view of the faradaic, double-layer capacitive, and mass-transport diffusion processes occurring within the layered bulk of the film. Faradaic redox reactions stemming from T_x_ groups of Ti_3_C_2_T_x_ flakes (OH^-^, H^+^, F^-^). Capacitive charge-transfer from Inner-and Outer Helmholtz Planes (IHP, OHP). Diffusion comes from the mass transport of larger ionic species.

Importantly, the diameter-dependent impedance trends observed here in this study have direct implications for recording noise and signal-to-noise ratio (SNR) when translated to in vivo applications. This relationship is fundamentally tied to the RMS thermal (Johnson) noise voltage (V_n,rms_) associated with the electrode impedance, which can be expressed as seen in Equation 4: ^72,83^

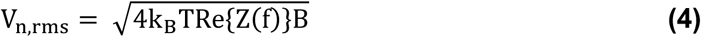

where k_B_ is the Boltzmann constant, T is the absolute temperature, Re{Z(f)} is the real component of impedance at frequency, f, and B represents the bandwidth of interest (typically 1, if f is a single frequency).^84^ Because V_n,rms_ scales with the square root of the real part of the impedance, Re{Z}, reductions in impedance at physiologically relevant frequencies are expected to lower the electrode noise floor and thereby improve recording quality. The resulting SNR can then be written as follows (**Equation 5**):

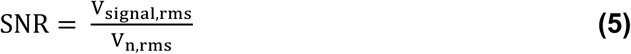

where V_signal,rms_ is the RMS amplitude of the signal of interest, and V_n,rms_ is the RMS amplitude of the noise signal, calculated above. Consistent with these relationships, and the EIS trends shown in **Figure 2**, when the multi-size Ti_3_C_2_T_x_ MXene microelectrodes were connected to a recording acquisition device, placed in a saline-bath, and subjected to an externally applied 250 µV, 1ms square voltage pulse – sent through the saline with an external stimulator – there was a clear and systematic increase in the measured SNR with increasing electrode diameter (**Figure S8**). Collectively, these relationships highlight why maintaining low, broadband impedance at microscale diameters is critical for high-fidelity electrophysiology: as electrode size decreases, impedance-driven thermal noise can become a dominant limitation even when amplifier noise is minimized. The relatively low impedance of Ti_3_C_2_T_x_ across frequencies, therefore, provides a materials-level pathway to suggest how to preserve recording quality as contact diameters shrink.

To extend these diameter-impedance relationships to practical device design considerations, we used the empirically fitted impedance-diameter and CIC scaling relationships to generate tentative, model-based estimates of the minimum electrode diameters required under representative recording and stimulation scenarios in different species (**Table S9**). These calculations are intended only as data-informed design guidelines and illustrative scaling projections rather than concrete in vivo limits. Accordingly, this table should be interpreted as a simplified framework for exploring tradeoffs between size, impedance, and charge delivery, and the predicted limits will require direct experimental validation under application-specific conditions.

Beyond these system-level performance considerations, it is also informative to examine how the intrinsic electrochemical properties of Ti_3_C_2_T*_x_* – particularly double-layer capacitance – scale with electrode geometry. Interestingly, even after normalizing the measured double-layer capacitance by the electrode geometric surface area, C_dl_ consistently increases as the contact diameter decreases. This trend likely arises because, as the electrode diameter decreases, a greater fraction of the total charge storage comes from ions accessing the internal volume of the spray-coated film rather than just the exposed surface; in other words, as the electrode footprint shrinks, film thickness and internal access pathways increasingly dominate the accessible charge storage per unit area. Practically, this suggests that, for device scaling, prioritizing thickness (and film layering that preserves ionic access) can partially offset the loss of geometric area and stabilize performance and CV-derived capacity as contacts approach the 10 µm - 100 µm regime.

From the voltage transient analysis, the charge injection capacity for both Ti_3_C_2_T*_x_* and Pt is relatively stable across all diameters, with only modest increases for the smallest contacts. Other work has noted edge-related scaling of CIC in larger planar electrodes, where current crowding at sharp electrode-encapsulation boundaries can elevate local charge density at the perimeter;^85^ however, in our benchtop voltage transient measurements, we did not observe pronounced edge effects across the 25 - 500 µm range.^34,79,82^ This muted diameter dependence is consistent with the specific device geometry used here, where, in our architecture, the exposed MXene aperture is bounded by a Parylene encapsulation overhang, and the MXene surface is recessed relative to the encapsulation edge due to the RIE patterning step.^86–88^ These features produce a non-planar, partially recessed perimeter that is analogous to recessed-electrode configurations previously shown to redistribute current density more uniformly across the electrode surface and suppress edge-localized charge amplification (**Figure S9**).^86–88^ Importantly, this does not preclude edge-profile-dependent differences *in vivo*, where current spread, tissue anisotropy, and encapsulation can amplify edge fields. Thus, the muted trend shown here *in vitro* may not generalize directly to biological stimulation conditions *in vivo*.^79,80^ Further, systematic variation of pulse width revealed a non-linear inverse relationship between maximum deliverable current and pulse duration, while the total charge per phase increased approximately linearly with pulse width. These relationships illustrate a fundamental trade-off between temporal precision and charge delivery: shorter pulses enable higher instantaneous current but less total charge transfer, whereas longer pulses increase charge storage at the expense of temporal resolution. Building upon these waveform-dependent findings, the subsequent analyses examine how film processing parameters – specifically deposition concentration and volume – affect interfacial charge transfer and the balance between recording and stimulation performance.

The two orthogonal processing parameters – dispersion concentration and spray-deposition volume – allowed us to separately evaluate the influence of surface roughness and film thickness on electrode behavior. When the dispersion concentration was increased from 1 mg mL^-1^ to 5 mg mL^-1^ while maintaining a fixed deposition volume of 50 mL, the resulting Ti_3_C_2_T*_x_* films exhibited increased surface roughness (root-mean-square roughness rising from ∼10 nm to ∼21 nm). Although this morphological change produced only minor variations in the overall impedance magnitude, the charge-transfer resistance decreased, and both the double-layer capacitance and cathodal charge storage capacity increased. These findings suggest that higher dispersion concentration enhances the number of electrochemically active sites and the accessible surface area within the film. Notably, similar to the impedance magnitude, the charge injection capacity remained largely unchanged, indicating that stimulation performance is comparatively insensitive to surface roughness within the tested range. When looking at dispersion volume, it was clear that increasing the deposited volume from 25 mL up to 100 mL at a fixed dispersion concentration of 3 mg mL^-1^ produced substantially thicker films (∼210 nm to ∼1,070 nm), which led to pronounced increases in double-layer capacitance and CSCc, consistent with greater volumetric ionic accessibility. Indeed, Nyquist plots for 200 µm Ti_3_C_2_T_x_ electrodes fabricated at these four deposition volumes show a systematic enlargement and leftward shift of the low-frequency semicircle with increasing film thickness, supporting its assignment within the equivalent circuit model to a thickness-dependent bulk ionic transport process throughout the layered MXene film (**Figure S10**). Despite this, the impedance magnitude and CICc values again remained relatively constant across thicknesses, reinforcing the idea that recording performance and stimulation safety limits depend primarily on the geometric surface area of the electrode rather than its bulk volume or surface roughness. It is worth noting, however, that based on these trends, the most advantageous film configuration would likely be one with both the highest volume and concentration, as this would minimize R_ct_, maximize C_dl_, and potentially increase long-term stability by forming a denser film and thicker barrier to oxidation.^89^ Consistent with this expectation, Ti_3_C_2_T_x_ microelectrodes spray-coated with 100 mL of 3 mg mL^-1^ that were stored under ambient conditions for 20 months maintained stable electrochemical behavior (validated with EIS and CV), indicating minimal degradation of impedance and charge storage capacity over time (**Figure S11, S12**). Fabrication limitations, however – such as spray-coater clogging at higher concentrations (> 5 mg mL^-1^) and a reduced yield of functional contacts due to delamination during the liftoff of the photomask – prevent an in-depth investigation of this optimal condition in this study. Ultimately, however, these results demonstrate that film thickness and porosity/roughness can be optimized to enhance the underlying charge-transfer mechanisms driving performance – manifested as reduced charge-transfer resistance, increased double-layer capacitance, and increased charge storage – while overall recording and stimulation performance remain predominantly governed by the electrochemically active surface area.

Finally, it is important to note that the benchtop measurements detailed in this study quantify intrinsic interfacial properties and recording and stimulation limits under controlled conditions and representative electrolytes, and because of this, there are several limitations that must be acknowledged. First, tissue coupling, encapsulation, physiological environment, and device morphology will introduce additional constraints *in vivo* (including possible edge-profile amplification and anisotropic current spread) that are not directly measured in this study.

Additionally, protein adsorption and the cerebrospinal fluid environment are expected to influence absolute impedance and charge-transfer behavior in vivo; therefore, the trends and scaling relationships reported here are intended to define comparative interfacial electrochemical limits and design tradeoffs rather than serve as direct predictors of chronic implanted performance. Separately, mechanical behavior under repeated bending and flexing represents a distinct device-level reliability question and is therefore outside the scope of the present work, but remains an important direction for future investigation. Within these bounds, however, the results from this work indicate that Ti_3_C_2_T*_x_* thin films combine exceptionally high interfacial capacitance with low charge-transfer resistance, enabling stable, low magnitude impedance across broadband frequency ranges and high CICc at microscale diameters without sacrificing safety. The processing variables investigated (thickness and roughness) further provide practical routes to stabilize recording performance during miniaturization while acknowledging that stimulation limits remain largely area-controlled. Further, while this study investigated performance variability within processing with spray-coating, it is important to acknowledge that there are several other processing deposition methods for fabricating MXene-based microelectrodes, including dip-coating, blade-coating, and spin-coating, that could affect the functional, morphological, and electrochemical performance (**Table S10**). Ultimately, however, by tuning the interplay between spray-coated volume, surface roughness, and electrode geometry, MXene-based electrodes can be optimized for scalable, high-density, high-fidelity neural interfacing technologies.

## CONCLUSIONS

In this work, we systematically investigated the electrochemical performance of Ti_3_C_2_T*_x_* MXene thin-film microelectrodes across a wide range of contact sizes (25 µm - 500 µm) and benchmarked them against thin-film platinum controls (**Table 1**). Using electrochemical impedance spectroscopy, cyclic voltammetry, and voltage transient analysis via multi-step chronopotentiometry, we demonstrated that Ti_3_C_2_T*_x_* electrodes maintain lower impedance, higher charge storage capacity, and greater charge injection capacity compared to platinum at matched geometries, including at the smallest contact sizes tested (25 μm). These results highlight the ability of MXene films to sustain efficient recording and stimulation performance when integrated into electrodes as their dimensions are miniaturized to tens of microns. Beyond these measurements, the equivalent circuit modeling further confirmed that Ti_3_C_2_T*_x_* microelectrodes exhibit behavior with strong capacitive contributions and exceptionally low charge-transfer resistance, consistent with efficient interface charge transfer for emerging neuroelectronics. This capacitive dominance, together with the material intrinsic conductivity and layered architecture, underpins the favorable scaling behavior observed across electrode sizes. Finally, we found that device fabrication parameters -- specifically MXene dispersion concentration and spray-coated volume -- significantly influence electrochemical outcomes. Together, this work establishes Ti_3_C_2_T*_x_* MXene as a versatile and scalable electrode material for next generation bioelectronic interfaces. Its ability to combine low broadband impedance, safe electrical stimulation, biocompatibility, flexibility, and compatibility with miniaturized, high-density architectures addresses critical challenges in the design of implantable electrode arrays.^46,60^ More broadly, this study provides a framework linking electrode geometry, material properties, clinically relevant stimulation pulse parameters (amplitude, pulse-width), and processing conditions to electrochemical performance, informing the rational development of bioelectronic systems for both basic research and clinical applications.

**Table 1.** Key electrochemical parameters of Ti_3_C_2_T*_x_* and sputtered Pt microelectrodes.

| | Diam.<br>($\mu\text{m}$) | 1 kHz $\vert Z\vert $<br>($\text{k}\Omega$) | 10 Hz $\vert Z\vert $<br>($\text{k}\Omega$) | $R_s$<br>($\text{k}\Omega$) | $R_{ct}$<br>($\text{k}\Omega$) | $C_{dl}$<br>($\mu\text{F cm}^{-2}$) | $\text{CSC}_c$<br>($\text{mC cm}^{-2}$) | $\text{Curr}_{max}$<br>($\mu\text{A}$) | $\text{CIC}_c$<br>($\mu\text{C cm}^{-2}$) |
| --- | --- | --- | --- | --- | --- | --- | --- | --- | --- |
| **$\text{Ti}_3\text{C}_2\text{T}_x$ MXene** | 500 | 1.9 | 4.7 | 0.5 | 39.9 | 3,820 | 20.8 | 1880 | 480 |
|  | 350 | 3.4 | 8.1 | 0.7 | 60.4 | 4,500 | 24.8 | 892 | 464 |
|  | 200 | 7.9 | 21.2 | 0.9 | 111 | 6,020 | 32.5 | 365 | 581 |
|  | 150 | 11.6 | 28.3 | 1.2 | 161 | 7,830 | 43.4 | 231 | 654 |
|  | 100 | 23.6 | 58.2 | 1.7 | 246 | 8,470 | 59.7 | 97.3 | 619 |
|  | 75 | 49.8 | 117 | 2.4 | 380 | 8,520 | 61.5 | 59.0 | 668 |
|  | 50 | 90.8 | 188 | 4.0 | 493 | 14,300 | 95.2 | 30.6 | 779 |
|  | 25 | 328 | 550 | 8.4 | 966 | 27,600 | 119.9 | 4.9 | 497 |
| **Sputtered Platinum** | 500 | 1.1 | 74.4 | 0.6 | 585 | 57.5 | 0.7 | 134 | 34.0 |
|  | 350 | 1.9 | 145 | 0.9 | 977 | 66.5 | 0.7 | 102 | 52.8 |
|  | 200 | 4.3 | 347 | 1.0 | 1,850 | 66.9 | 0.8 | 67.6 | 108 |
|  | 150 | 6.7 | 529 | 1.6 | 2,580 | 64.7 | 0.9 | 33.6 | 95.0 |
|  | 100 | 13.7 | 1,100 | 2.5 | 4,390 | 83.0 | 0.9 | 22.9 | 146 |
|  | 75 | 18.5 | 1,390 | 3.3 | 4,840 | 102 | 1.0 | 15.0 | 170 |
|  | 50 | 49.0 | 2,670 | 5.2 | 10,200 | 145 | 1.9 | 5.6 | 144 |
|  | 25 | 140 | 4,370 | 6.1 | 11,200 | 213 | 4.3 | 1.3 | 141 |

## METHODS

### Ti_3_C_2_T_x_ MXene Synthesis

Suspensions of Ti_3_C_2_T*_x_* MXene were produced and provided by MuRata Manufacturing Co., Ltd. Large Ti_3_C_2_T_x_ flakes (∼ 1.3 µm lateral size) with a Zeta potential of approximately -20 mV were synthesized using the minimally intensive layer delamination (MILD) method, where the Al is removed from the Ti_3_AlC_2_ powder (MAX phase) with a combination of 12 M LiF and 9 M HCl.^45,90–92^ After synthesis and prior to spray-coating, the Ti_3_C_2_T*_x_* MXene suspensions were stored in a large glass vial at 4 °C. The stock 35.6 mg mL^-1^ Ti_3_C_2_T_x_ MXene suspensions were diluted with deionized water to 3 mg mL^-1^ for the fabrication of the Ti_3_C_2_T*_x_* thin-film microelectrodes. A subset of the stock Ti_3_C_2_T*_x_* was diluted to 1 mg mL^-1^ and 5 mg mL^-1^ for the investigation of the effects of Ti_3_C_2_T*_x_* concentration on the performance of the microelectrodes.

### Thin-film Ti_3_C_2_T_x_ MXene Microelectrode Fabrication

Thin-film microelectrodes were fabricated using protocols previously established. First, 3.5 g of Parylene-C was deposited via chemical vapor deposition to form a 4-µm thick substrate layer on a 3” mechanical-grade silicon wafer. Then, negative photoresist (NR7-3000p) was spun at 3000 rpm onto the wafer which photolithographically patterned the backend channels which were made by electron beam deposition of 10 nm Ti followed by 60 nm Au. Residual metal and photoresist were lifted-off using an organic solvent (KL Remover 1000). A second negative photoresist layer was spun onto the wafer and photolithographically defined the pattern for the Ti_3_C_2_T*_x_* contacts. Prior to spray-coating of the Ti_3_C_2_T*_x_*, a 3-minute reactive ion etch (RIE) using ozone plasma primed the surface of the Parylene, where the negative photoresist outlined the contact locations. The Ti_3_C_2_T_x_ MXene dispersion (3 mg mL^-1^) was spray-coated onto the wafer using a handheld airbrush operated at 30 PSI. The dispersion was sprayed perpendicular to the wafer surface from a distance of approximately 2 ft, with deposition performed while the wafer rested on a 130 °C hot plate to promote rapid evaporation of the aqueous solvent upon contact. During spraying, the wafer was rotated in 90° increments at regular intervals to ensure uniform coverage and homogeneous material distribution across the exposed contact regions. Following deposition, the wafer was placed in an 80 °C oven for 30 min to dry any residual Ti_3_C_2_T_x_, after which the remaining MXene and photoresist were lifted off using an organic solvent bath at 70 °C. A final top-passivation parylene-C layer was then deposited – again using 3.5 g parylene-C pellets – to completely seal the microelectrodes. A final top-passivation parylene-C layer was then deposited – again using 3.5 g parylene-C pellets – to completely seal the microelectrodes. A final negative photoresist layer defined the outline of each device as well as the electrode contacts and backend connection pads. A 100 nm-thick Al layer was then electron-beam deposited and excess was lifted off, leaving the contact and backend pads exposed without any aluminum. A final 15-minute oxygen RIE etch was performed, etching away the parylene-C that covered the contacts, backend pads, and complete devices (i.e. the non-Al coated regions). Finally, the wafer was dipped in an aluminum etchant and complete devices were lifted off the wafer immediately prior to electrochemical characterization. Prior to use, all devices were stored in a N_2_-purged desiccator.

### Thin-film Sputtered platinum Microelectrode Fabrication

Pt microelectrodes were fabricated using an identical protocol as detailed above. The key difference was that after the second negative photoresist was spun on and photopatterned, without the need of ozone priming of the parylene surface, the wafer was placed in a magnetic sputterer and 10 nm Ti and 100 nm platinum was sputtered onto the wafer. Residual Pt and photoresist were then lifted off using an organic solvent and the wafer then resumed the fabrication protocol described above.

### Optical Microscopy

Optical microscopy images of the Ti_3_C_2_T*_x_* microelectrodes were acquired using 200x magnification on a Keyence VHX6000 digital microscope. 3D depth composition was performed during the imaging acquisition to resolve the image at multiple depths.

### Scanning Electron Microscopy

SEM images of the Ti_3_C_2_T*_x_* and Pt microelectrodes were collected using a FEI Quanta 600 FE-SEM machine. All samples were grounded to the system using a combination of copper and carbon tape. Image acquisition of the Ti_3_C_2_T*_x_* and Pt contacts was collected under high vacuum at approximately 6x10^-4^ Pa. The spot size was held to 3, and the voltage was set to 10 keV. Image magnification ranged from 100x - 2000x.

### EDX Spectroscopy

EDX spectra were collected on the same sample images that were acquired during SEM imaging, but the spot size was increased to 5 to increase the counts per second acquired by the sensor. Data collection was continued until an adequate signal-to-noise ratio was obtained, generally corresponding to at least ∼40,000 accumulated counts, which provides ∼±1% precision in elemental quantification. These standardized acquisition parameters yielded high-quality, reproducible EDX spectra across all microelectrode samples.

### Atomic Force Microscopy

Atomic Force Microscopy surface mapping was collected using a Bruker Dimension Icon AFM machine in tapping mode in air. A Silicon tip cantilever (*f_o_* = 300 kHz) was used to characterize the sample. Samples measured were 500 µm diameter DIW printed Ti_3_C_2_T*_x_* electrode contacts from a subset of the device used in the intraoperative experiment. The scan size was 10 μm with an aspect ratio of 1 and was collected at a scan rate of 1 Hz. All data and mean surface roughness calculations were analyzed in NanoScope Analysis software (Bruker).

### X-Ray Photoelectron Spectroscopy

XPS was acquired using a PHI VersaProbe 5000 instrument (Physical Electronics) with a 25 W monochromatic Al-Kα (1486.6 eV) X-ray source targeting a 200 μm spot. Charge neutralization was acquired through a dual-beam setup using low-energy Ar+ ions and low-energy electrons at 1 eV and 200 μA. Survey spectra and High-resolution spectra from the Ti 2p and C 1s core-level region were collected using a pass-energy/energy-resolution of 117.4 / 0.5 and 23.5 / 0.05 eV, respectively. The binding energy scales were calibrated by adjusting the C-C / C-H component from adventitious carbon in the C 1s spectra to 284.8 eV, where the required shift was < 3.3 eV for all samples. Finally, quantification and peak fitting were conducted using the CasaXPS software.

### Raman Spectroscopy

A Renishaw inVia Raman spectrometer (Gloucestershire, UK) instrument was used for all measurements. The acquisition time for the measurements was 10 seconds using a 63x objective (NA = 0.7). The excitation intensity using the diode laser (785 nm) was 10.5 mW, with a 10% power modulation. All measurements were conducted by raster scanning from 100-3200 cm^−1^.

### X-Ray Diffraction Spectroscopy

A Rigaku MiniFlex benchtop X-ray diffractometer was used for X-ray diffraction (XRD) measurements (Rigaku Co. Ltd.). The source was Cu Kα (λ = 0.1542 nm), and spectra were acquired at 40 kV, 15 mA, for Bragg angles from 3°–60° at a rate of 14° min–1 with a step of 0.02°. After collection, spectra were fitted using CrystalDiffract to identify peak locations.

### Three-Cell Electrochemical Setup

All electrochemical experiments were collected using a Gamry Reference 600 potentiostat. Measurements were conducted at room temperature in 0.01 M (1x) phosphate-buffered saline (PBS, pH 7.4). The measurements were collected in a standard three-electrode electrochemical setup, with the thin-film Ti_3_C_2_T*_x_* MXene or platinum microelectrode contact as the working electrode, a long graphite rod (Bio-Rad Laboratories, Inc.) as the counter electrode, and a single-frit Ag/AgCl electrode filled with saturated potassium chloride (KCl, 3 M) as the reference electrode. For all electrochemical experiments, voltage referencing was performed at 0 V vs the Ag/AgCl reference electrode. We characterized the electrochemical interface of Ti_3_C_2_T*_x_* and platinum microelectrodes by measuring the impedance modulus (|Z|), charge storage capacity (CSC), and charge injection capacity (CIC) of all contacts. The impedance was acquired using electrochemical impedance spectroscopy (EIS), voltage limits (potential window) and CSC were determined from cyclic voltammetry (CV), and maximum stimulation currents and CIC were calculated from voltage transient (VT, a.k.a. multi-step chronopotentiometry) measurements.

### Electrochemical Impedance Spectroscopy

Electrochemical Impedance Spectroscopy (EIS) curves were collected by applying a 10 mV_rms_ sinusoidal AC voltage at decreasing frequencies from 100 kHz down to 1 Hz. Equivalent circuit modeling was performed using an open-source Python package (Impedance.py).^93^ Once the equivalent circuit model was established, impedance spectra from each microelectrode were individually fit using the impedance.py package, and the resulting parameter values were averaged across electrodes for all reported results.

### Cyclic Voltammetry

Cyclic voltammetry (CV) measurements were also collected using the Gamry Reference 600 potentiostat in the same three-cell setup as described above. The anodic and cathodic voltage limits for the Ti_3_C_2_T_x_ microelectrodes were measured by sequentially increasing the cathodic and anodic potential limit until the voltammogram skewed towards O_2_ or H_2_ formation, where the limits closely approximate values from previously published work, and were held fixed at -1.55 V for the cathodic limit and 0.35 V for the anodic limit.^55^ The potential window limits for the Pt microelectrodes have been established in multiple prior works and were held fixed at -0.6 V and 0.8 V.^13,23,76^ CVs were collected at 500 mV/s. Measurements started at 0 volts and scanned in the positive direction first. Each microelectrode was subjected to 10 cycles of CV scans at a resolution of 2 mV, and analysis was conducted on the 10^th^ and final scan. Additional CVs were collected for each electrode at a range of scan rates from 50 mV/s up to 500 mV/s over a limited (and matched, -0.3 V to 0.3 V) voltage range material to assess and compare specific capacitance. Cathodal charge storage capacity (CSCc) was calculated for each electrode from the 10^th^ scan of each contact size for each material as the time integral of the negative current during a full CV at 500 mV/s.

### Voltage Transients (multi-step chronopotentiometry)

Voltage transient measurements were cathodic-leading and adjusted to a baseline of 0 volts after being collected for analysis. All stimulation pulses were biphasic and charge-balanced. The anodic current was twice the duration at half the current amplitude as the cathodic pulse to maintain charge balance and remain below the low anodic potential window limit of Ti_3_C_2_T*_x_* (0.35 V). Each electrode was subjected to a series of 15-25 stimulation pulses at increasing current amplitudes around their expected maximum current limit. For the charge injection capacity (CIC) calculations, E_mc_ was determined as the instantaneous voltage 10 µs after the end of the cathodic current pulse. Maximum current amplitude and resulting charge injection capacity (CIC) were then calculated using the largest applied current amplitude where E_mc_ did not exceed the cathodic potential window limits. For this study, none of the current pulses exceeded the anodic potential window limit, but rather only passed the cathodic potential window limit due to the waveform asymmetry. Because of this, the reported CIC was calculated only using the cathodic current amplitude and was based on the charge injected per phase. CIC was calculated using **Equation 6**, where PW is the pulse width (for most analyses, this was 500 µs), I_max_ is the maximum sub-potential window limit current amplitude, and A is the electrode’s surface area. Stimulation pulse-widths were varied from 100 µs cathodic pulses up to 1500 µs cathodic pulses, and the resulting CICs were measured as a function of not only contact size but also stimulation pulse-width. The clear trend was that CIC increased with increasing pulse-width, at the cost of decreasing the maximum current amplitude.

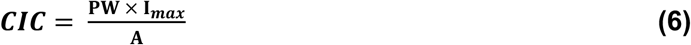

### Profilometry Measurements

Film thickness was characterized using a KLA Tencor P7 profilometer. Measurements were conducted with an applied stylus force of 1 mg, scan speed of 20 µm s^-1^ and sampling rate of 100 Hz. The instrument provided a vertical range of 327 µm with a vertical resolution of 0.1953 Å. Each scan was 2mm in length. For analysis, the scan was leveled relative to the bare glass substrate using the profilometers level function, and the film height was averaged over a representative plateau region. For concentration-dependent measurements, MXene films were spray-coated from 1, 3, and 5 mg mL^-1^ Ti_3_C_2_T*_x_* dispersions (50 mL total volume) onto three glass slides per condition. A dull blade was used to gently remove a portion of the coating to expose the underlying glass, which served as the reference height for leveling. For volume-dependent measurements, profilometry was performed directly on representative device contacts. A total of fifteen measurements were collected across all tested conditions.

### Shelf Life Stability

To evaluate long-term ambient storage stability (“shelf-life”), a subset of Ti_3_C_2_T_x_ MXene microelectrode arrays were stored under ambient conditions (room temperature, atmospheric air, no environmental sealing or desiccation) for approximately 20 months following fabrication. No additional encapsulation or protective coatings were applied beyond those described in the fabrication methods above. After storage, electrochemical performance was reassessed and compared to that of freshly fabricated devices from the same fabrication process. Electrochemical impedance spectroscopy (EIS) and cyclic voltammetry (CV) were performed following the same procedures, electrolyte, electrode configuration, and instrumentation described above for as-fabricated devices to ensure direct comparability. For EIS, impedance magnitude and phase spectra were collected across the same frequency range and with the same perturbation amplitude as described previously, with impedance values at 1 kHz and 10 Hz extracted to evaluate changes in interfacial behavior. Measurements were obtained from multiple electrodes per diameter for both stored (n = 4 per size) and as-fabricated (n = 34 per size) devices. For the CVs, voltammograms were recorded over the same potential window (-1.55 V to +0.35 V vs. Ag/AgCl) and scan rate used for freshly fabricated electrodes. Charge stored and charge storage capacity (CSC) were calculated using the same integration and analysis procedures described above. CV curves and derived charge metrics from stored devices were directly compared to those from as-fabricated electrodes of matching diameters.

### Equivalent Circuit Model Error Analysis

EIS spectra were analyzed using the open-source “impedance” Python package (impedance.py). Complex impedance data Z(ω)=Z′(ω)+jZ′′(ω) were imported as a function of frequency and assessed for compliance with the Kramers-Kronig (KK) constraints using the linear KK (Lin–KK) method (impedance.validation.linKK) Briefly, Lin–KK performs a least-squares reconstruction of the measured impedance spectrum using a flexible KK-compliant impedance representation (i.e., enforcing causality and linear time-invariant behavior) without assuming a specific equivalent-circuit topology. The algorithm returns the KK-consistent reconstructed spectrum Z_LinKK_(ω) as well as frequency-resolved residuals for the real and imaginary components. These residuals were computed as the difference between the measured and reconstructed impedance and were expressed as normalized (percent) errors relative to the complex impedance magnitude, i.e**., Δ_Re_(ω)=Re(Z_meas_−Z_fit_)/∣Z_meas_∣** and **Δ_Im_(ω)=Im(Z_meas_−Z_fit_) /∣Z_meas_|**. This normalization by the magnitude of the measured complex impedance at each frequency (|Z_meas_|) expresses residuals as relative (percent) error and enables comparison across electrodes of different sizes and absolute impedance values. To quantify model performance, equivalent-circuit parameters were fit to the measured data (complex-valued fit), and the same normalized residual definitions were applied to compute Δ_Re_(**ω**) and Δ_Im_(**ω**) for the circuit fit. Root-mean-square (RMS) percent error was calculated separately for the real and imaginary components as RMS_Re_=⟨(100 * Δ_Re_)^2^⟩ and RMS_Im_=⟨(100 * Δ_Im_)^2^⟩, where ⟨⋅⟩ denotes averaging across all frequency points over the KK-consistent frequency range (<10 kHz) to provide an error metric restricted to the linear/KK-supported regime.

### In-Vitro Signal-to-Noise Ratio (SNR) Calculations

To experimentally quantify the relationship between contact size and recording signal-to-noise ratio (SNR), we performed an in vitro saline-bath stimulation experiment using multi-size Ti_3_C_2_T_x_ MXene microelectrode arrays. Three independently fabricated devices were tested (n = 3 devices), each containing 16 total recording channels comprising eight contact sizes, with two replicate electrodes per size (n = 2 contacts per size per device; total n = 6 electrodes per contact size across devices). Devices were submerged individually in a saline bath contained in a small Petri dish and connected to a Ripple Grapevine acquisition processor for real-time multichannel recording. To generate a controlled, repeatable evoked response in the electrolyte, a pair of stimulation electrodes (cathode and anode) connected to a Keithley stimulus source were placed in the same saline bath at fixed positions relative to the recording array. A series of ten stimulation pulses (n = 10 pulses per device) were delivered as square-wave voltage steps (250 µV amplitude, 1 ms duration), applied one pulse at a time with several seconds between pulses to allow the recording baseline to stabilize between events. During stimulation, all 16 channels were recorded simultaneously and saved in NS5 format for offline analysis. SNR analysis was performed using custom Python scripts that imported Ripple NS5 recordings, identified stimulation events, and quantified electrode-wise evoked response amplitude relative to baseline noise. For each recording channel and each of the 10 stimulation pulses, a 250 ms pre-stimulus time window was used to characterize the baseline noise (RMS of the voltage trace), and a 250 ms stimulus response window was used to quantify the signal magnitude as the peak amplitude relative to baseline. SNR was then computed for each pulse and channel as the ratio of peak amplitude to baseline noise (reported in dB), and values were subsequently aggregated across stimulation pulses and replicate electrodes. Final SNR values for each contact diameter represent the mean SNR across all 10 stimulation pulses and electrodes of that size (n = 6 electrodes), enabling direct comparison of SNR trends across contact sizes under identical stimulation and recording conditions.

## ASSOCIATED CONTENT

### Supporting Information

#### Data Availability

The data supporting the findings of this study are available upon reasonable request from the authors.

## AUTHOR INFORMATION

Corresponding Author *

## Author Contributions

S.R.A. and F.V. initiated the project. R.G. provided assistance in calculating CSC from the cyclic voltammograms. R.D. collected all profilometry measurements. D.H. collected all Atomic Force Microscopy measurements on the MXene and Platinum electrode contacts. N.V.A. and M.S.B. provided additional supervision and technical feedback throughout the project. S.R.A. wrote the manuscript with contributions from all other authors. F.V. supervised the project.

## Conflict of Interest Statement

F.V. is a co-inventor on the US patent: US11925466B2 “Implantable devices using 2D metal carbides and nitrides (MXenes)”.

## ACKNOWLEDGEMENTS

This work was supported by the National Science Foundation (CAREER Award no. 2339748 to F.V.), and the National Institutes of Health (R01NS123054 to F.V.; R01NS065395 to M.S.B.; T32NS091006 and F30EY037143 to R.D.). Microfabrication was conducted in the Singh Center for Nanotechnology, which is supported by NSF NNCI-2025608. We acknowledge Murata Manufacturing Co. Ltd. for providing the Ti_3_C_2_T*_x_* MXene used in the study.

---

**TABLE OF CONTENTS.**
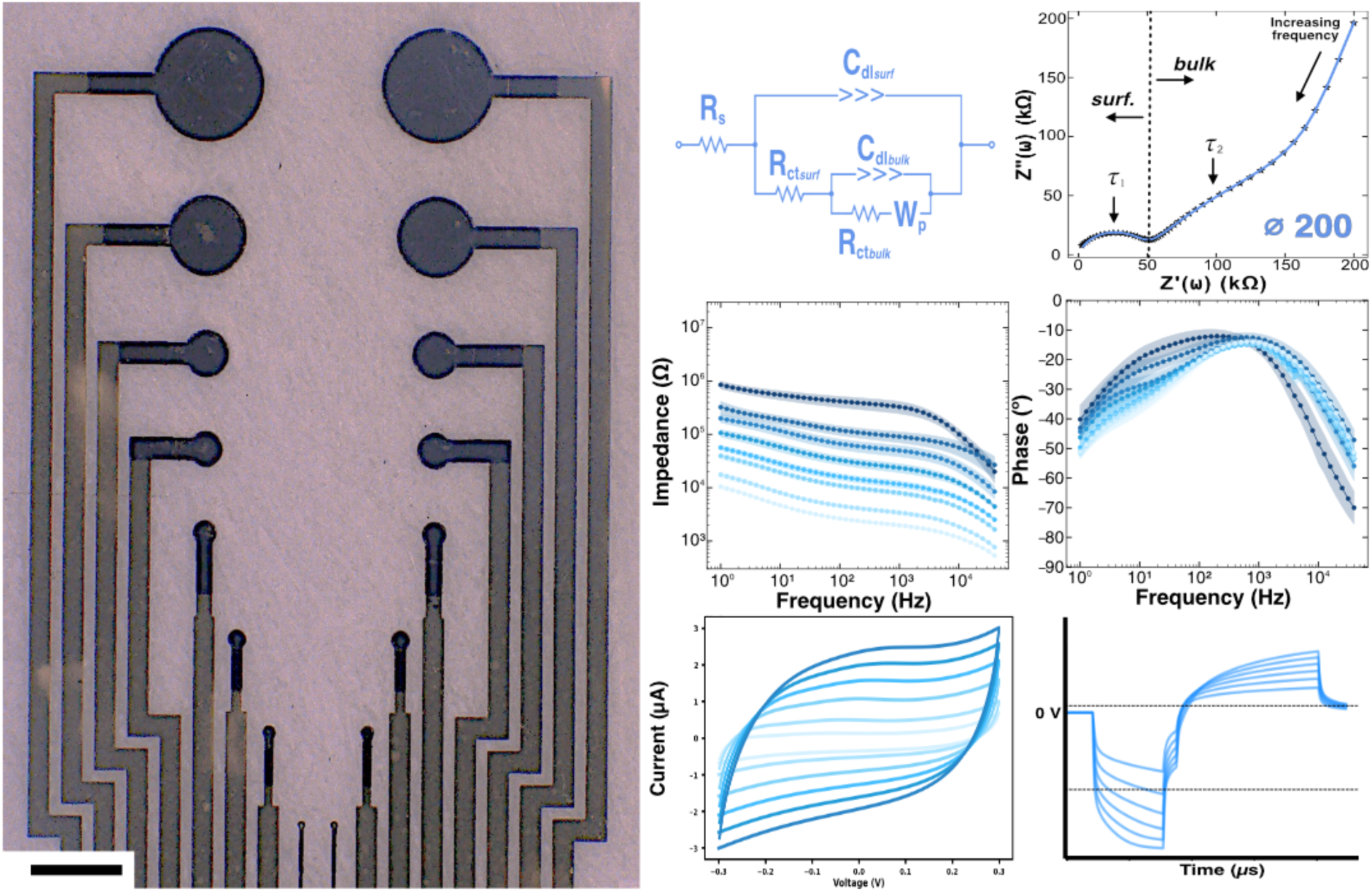
This study evaluates Ti_3_C_2_Tx MXene-based microelectrodes across a range of diameters (25 - 00μm) and elucidates their recording and stimulation capabilities for bioelectronic applications. Several electrochemical measurements - supported further by equivalent-circuit modeling - revealed enhanced recording and stimulation capabilities of the MXene microelectrodes over size-matched Pt microelectrodes, attributed predominantly to reduced charge-transfer resistance and increased double-layer capacitance.

## Notes

### Competing Interest Statement

The authors have declared no competing interest.

