## Supporting Information for "Size Scaling of the Electrochemical Performance of Ti_3_C_2_T*_x_* MXene Microelectrode Arrays for Electrophysiological Recording and Stimulation"

### SUPPLEMENTARY FIGURES

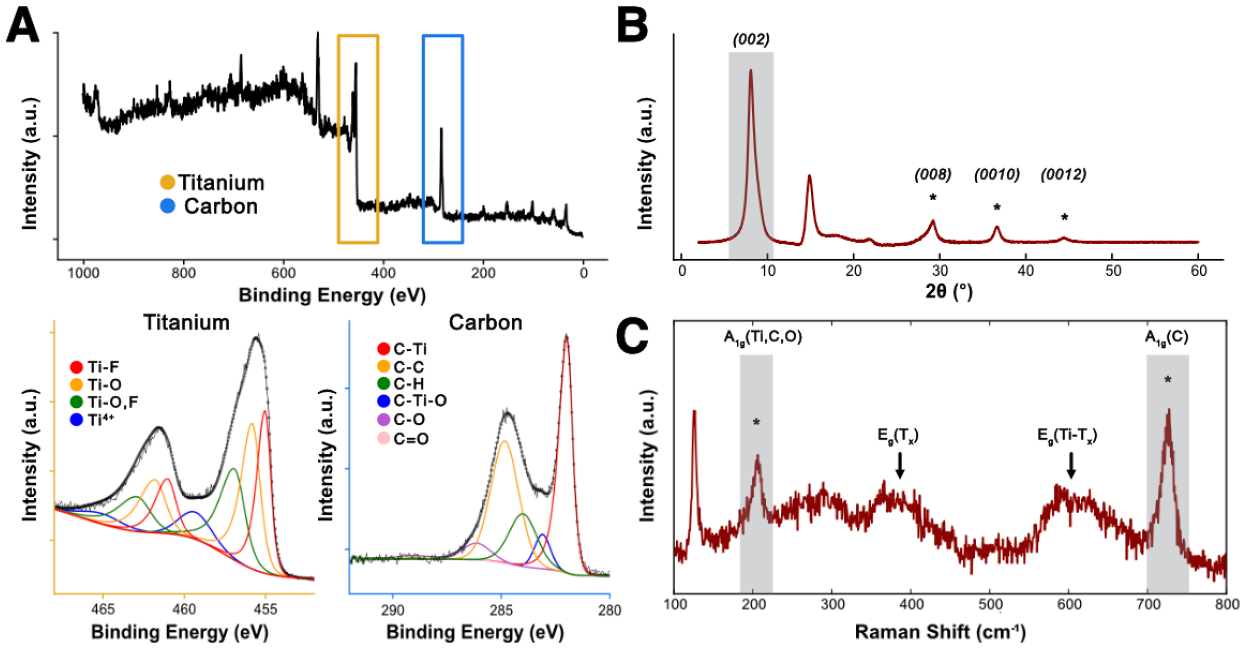


**Figure S1. Structural analysis of MILD-synthesized $Ti_3C_2T_x$ MXene. (A)** *Top:* XPS Survey Spectra of $Ti_3C_2T_x$ MXene, highlighting Titanium and Carbon peaks. *Bottom Left:* Core-level Ti *2p* spectra of $Ti_3C_2T_x$ MXene. *Bottom Right:* Core-level C *1s* spectra of $Ti_3C_2T_x$ MXene **(B)** XRD spectra of $Ti_3C_2T_x$ MXene, highlighting characteristic 002 peak. **(C)** Raman spectra of $Ti_3C_2T_x$ MXene, highlighting characteristic out-of-plane $A_{1g}$ peaks as well as the in-plane $E_g$ peaks.

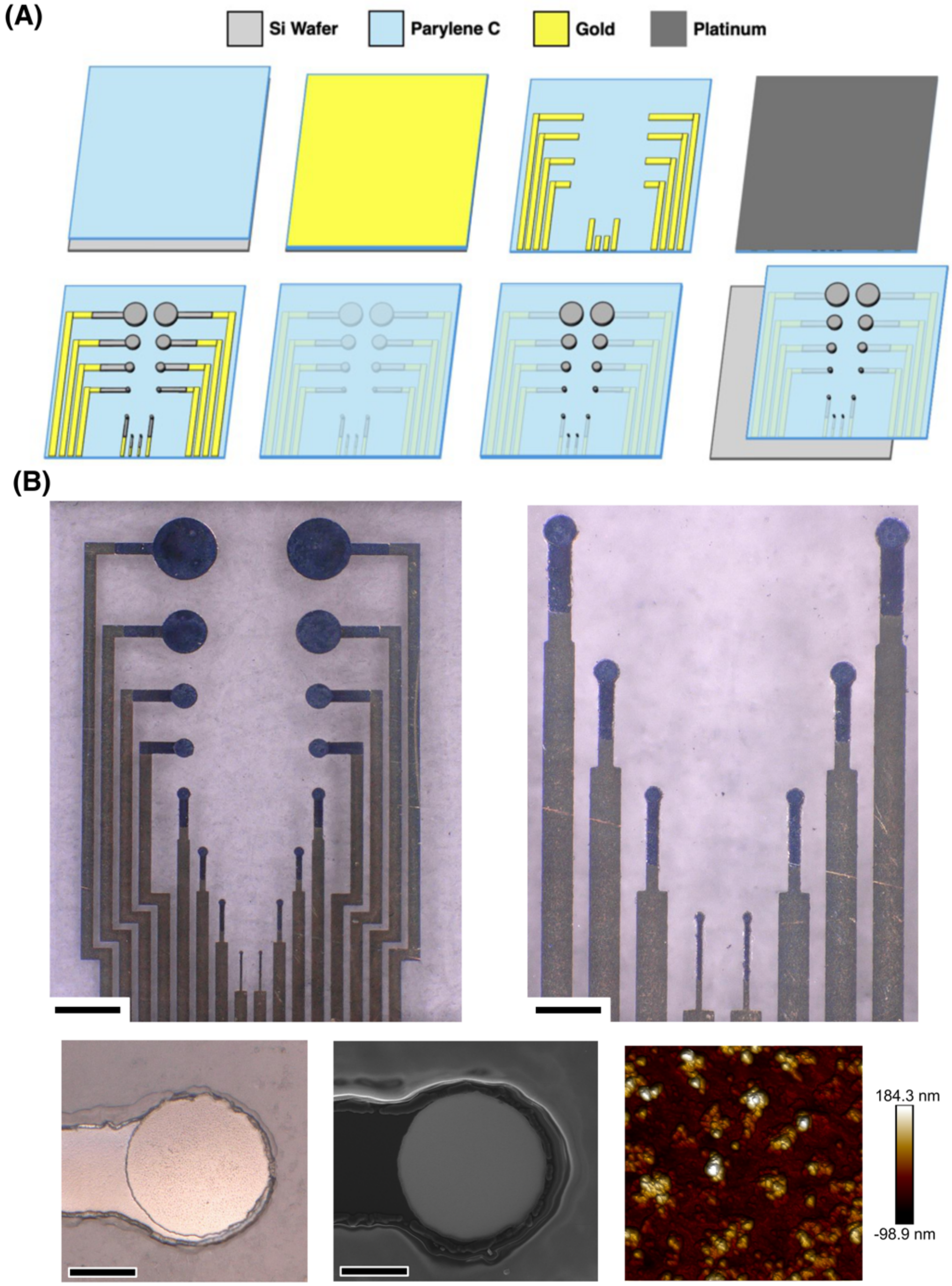


**Figure S2. Fabrication schematic and images of Pt microelectrodes. (A)** Schematic of the fabrication flow. **(B)** *Top:* Optical image of Pt microelectrodes (scale bar 500 μm) with zoomed-in insets of 100 μm, 75 μm, 50 μm, and 25 μm contacts. Scale bar 150 μm. *Bottom.* From left to right: optical, SEM, and AFM images of 100 μm Pt contacts. Scale bar 30 μm.

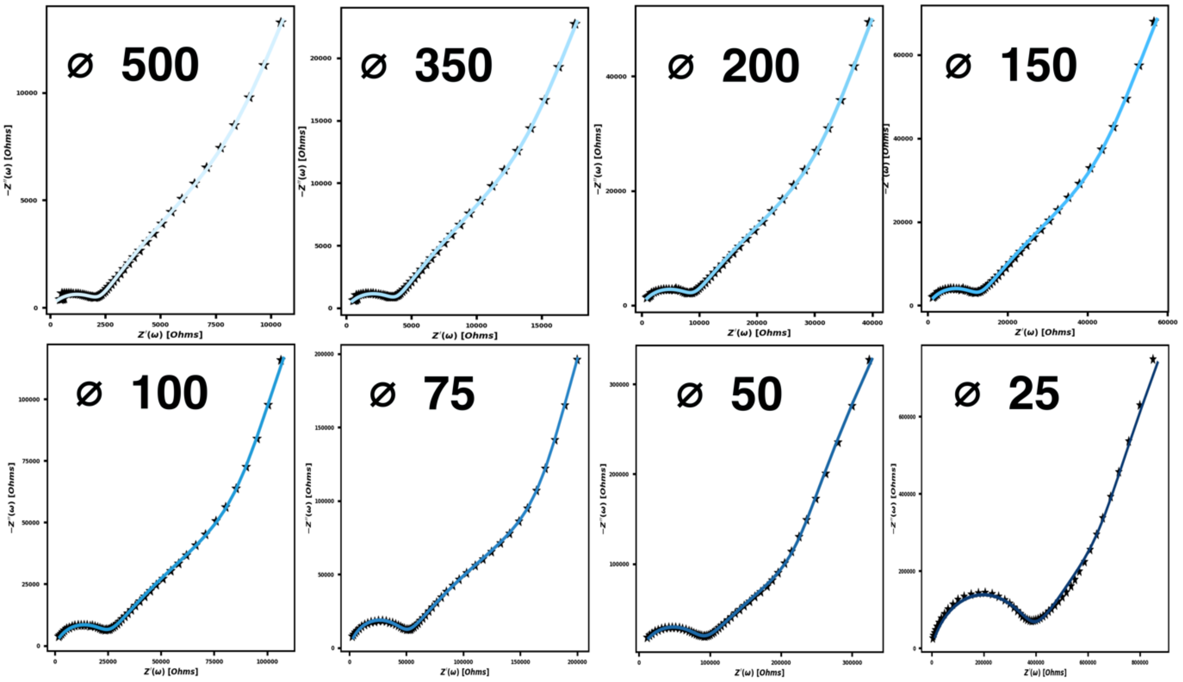


**Figure S3. $Ti_3C_2T_x$ microelectrodes: Nyquist plots and equivalent circuit model fittings for all contact diameters.**

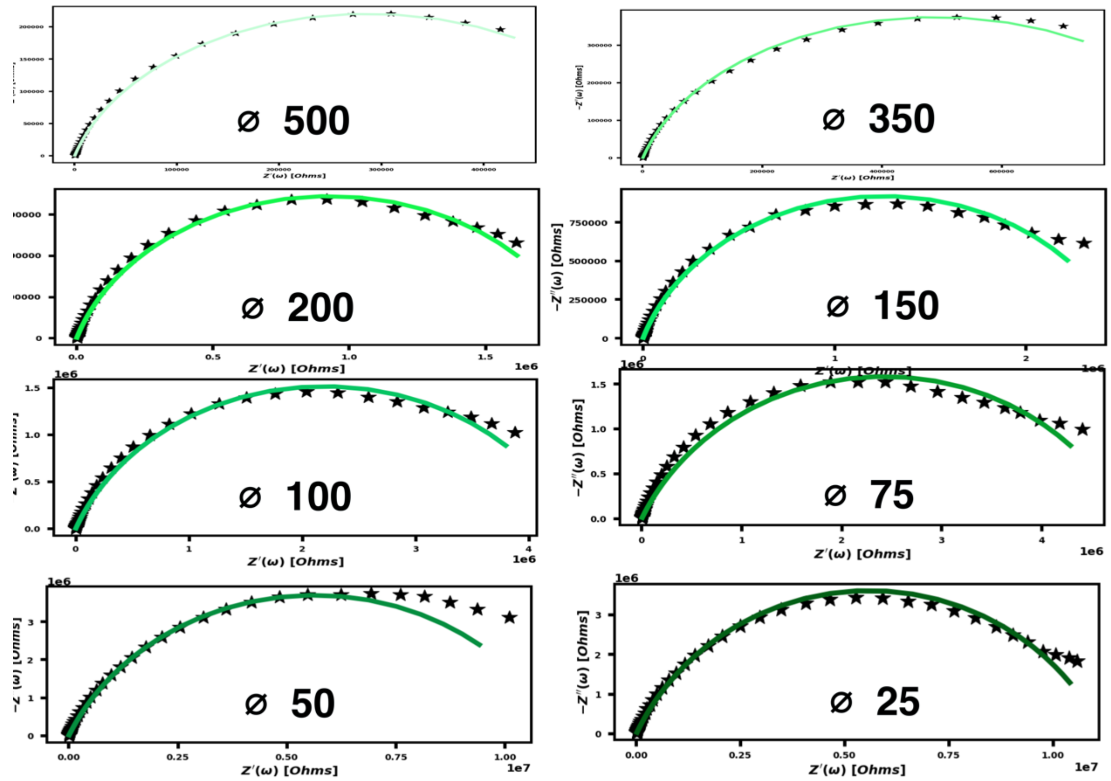


**Figure S4. Pt microelectrodes: Nyquist plots and equivalent circuit model fittings for all contact diameters.**

A

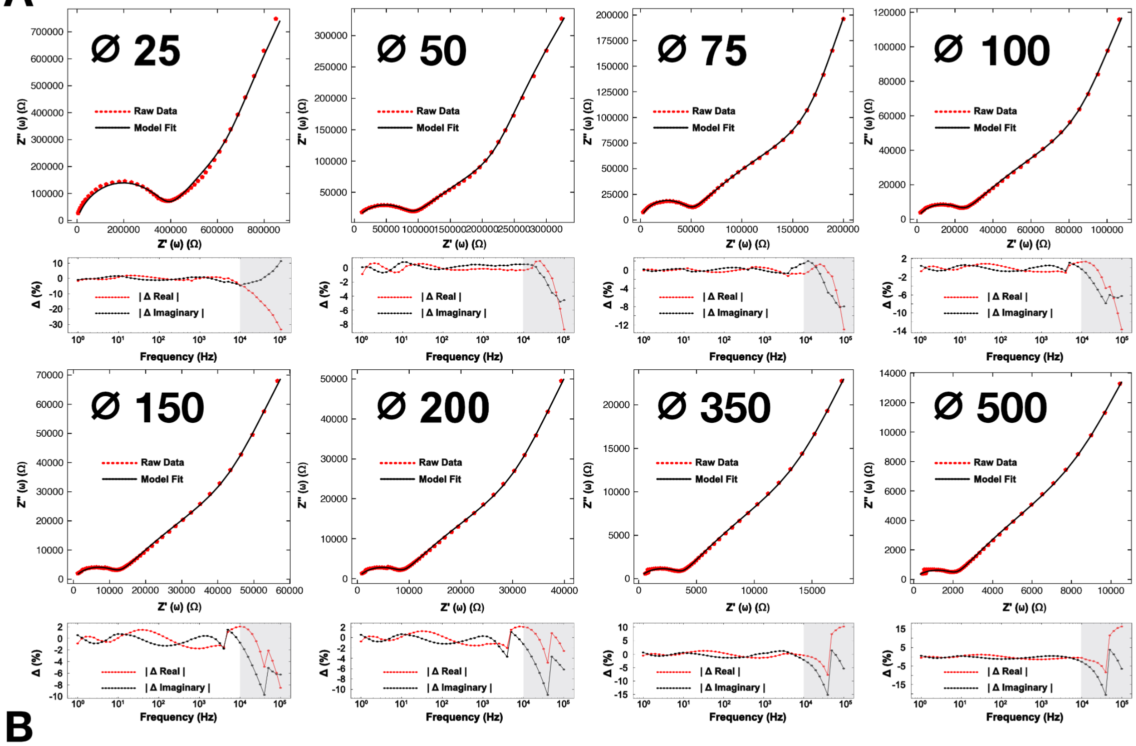


B

| Electrode Diameter | 25 µm | 50 µm | 75 µm | 100 µm | 150 µm | 200 µm | 350 µm | 500 µm |
|---|---|---|---|---|---|---|---|---|
| RMS Error$_{Real}$ (%) | 1.24 | 0.38 | 0.45 | 0.68 | 1.14 | 1.11 | 0.77 | 0.82 |
| RMS % Error$_{Imaginary}$ (%) | 1.39 | 0.41 | 0.47 | 0.51 | 0.77 | 1.00 | 0.85 | 0.80 |

**Figure S5. Error Analysis and Model Fit Residuals**. **(A)** For each contact diameter, *(Top)* Raw and fitted Nyquist plot, (*Bottom)* Real and imaginary model fit residuals across all frequencies, highlighting low error for all frequencies that satisfy KK-linearity consistency checks (1 Hz – 10 kHz). Grey-highlighted regions in residuals represent content that did not meet a KK consistency check within 2.5%. **(B)** RMS error for real and imaginary components of residuals within the KK-linear region, expressed as RMS percent error, where residuals at each frequency are normalized by the magnitude of the measured impedance $|Z_{meas}|$ for each electrode diameter.

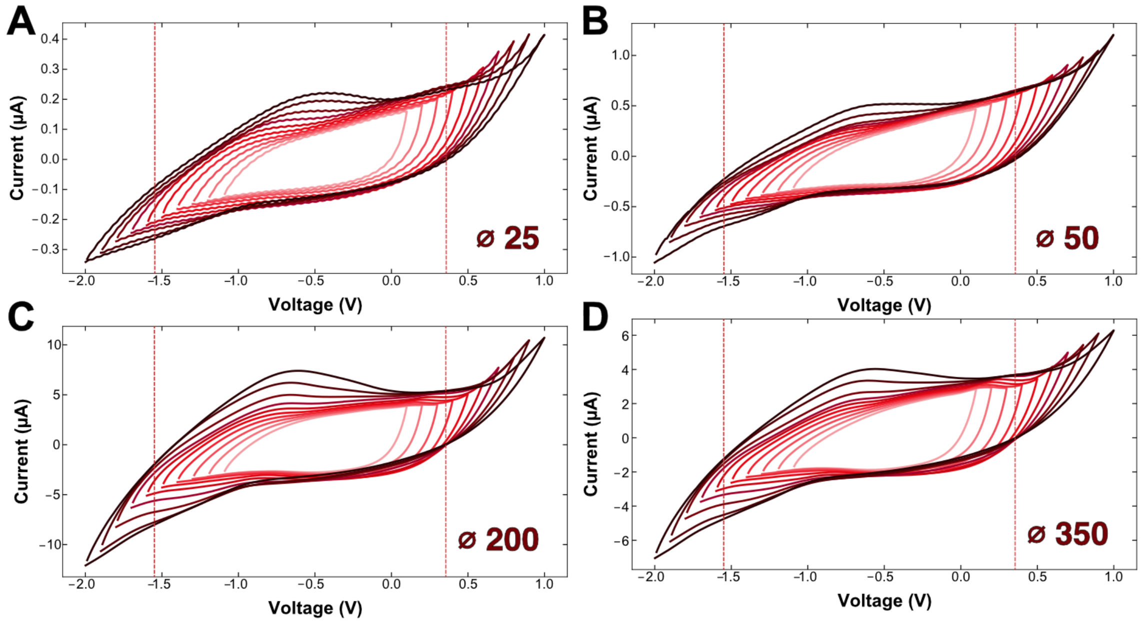


**Figure S6. Water window determination for $Ti_3C_2T_x$ MXene thin-film microelectrodes. (A-D)** Expanding potential windows delineating anodic and cathodic voltage limits for $Ti_3C_2T_x$ MXene thin-film microelectrodes at diameters of **(A)** 25 µm, **(B)** 50 µm, **(C)** 200 µm, and **(D)** 350 µm.

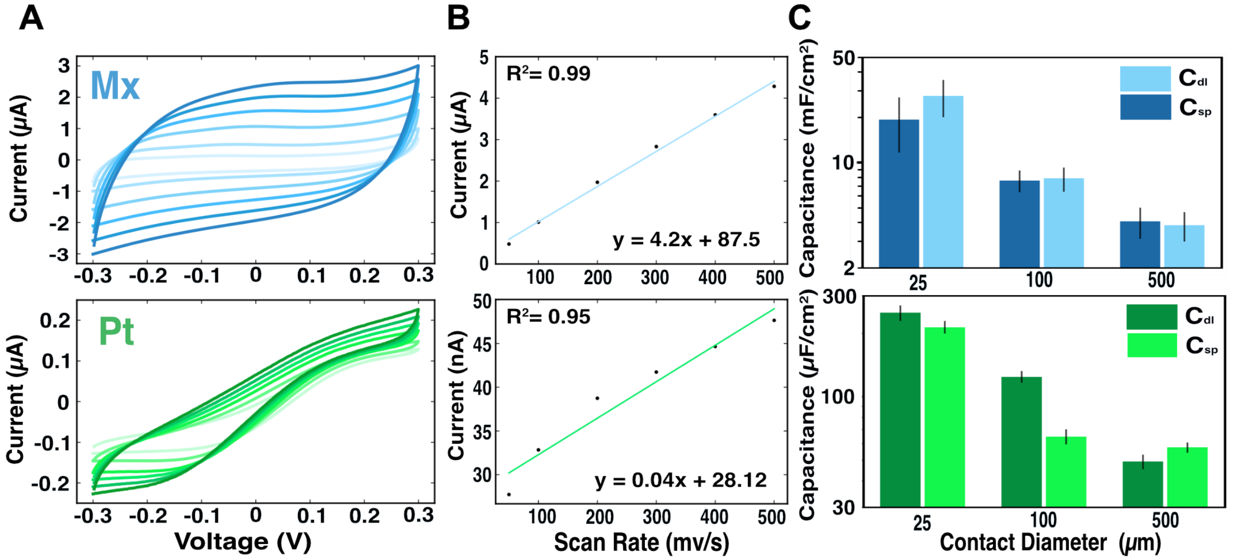


**Figure S7. Specific capacitance from CVs vs $C_{dl}$ from the equivalent circuit model. (A)** Cyclic voltammograms at varying scan rates (50 mV $s^{-1}$, 100 mV $s^{-1}$, 200 mV $s^{-1}$, 300 mV $s^{-1}$, 400 mV $s^{-1}$, 500 mV $s^{-1}$) for 350 µm microelectrodes. *Top:* $Ti_3C_2T_x$, *Bottom:* Pt. **(B)** Current vs. scan rate for *Top:* $Ti_3C_2T_x$ and *Bottom:* Pt microelectrodes. **(C)** Comparison of the capacitance calculated from **(B)** and from the equivalent circuit model for *Top:* $Ti_3C_2T_x$, *Bottom:* Pt microelectrodes.

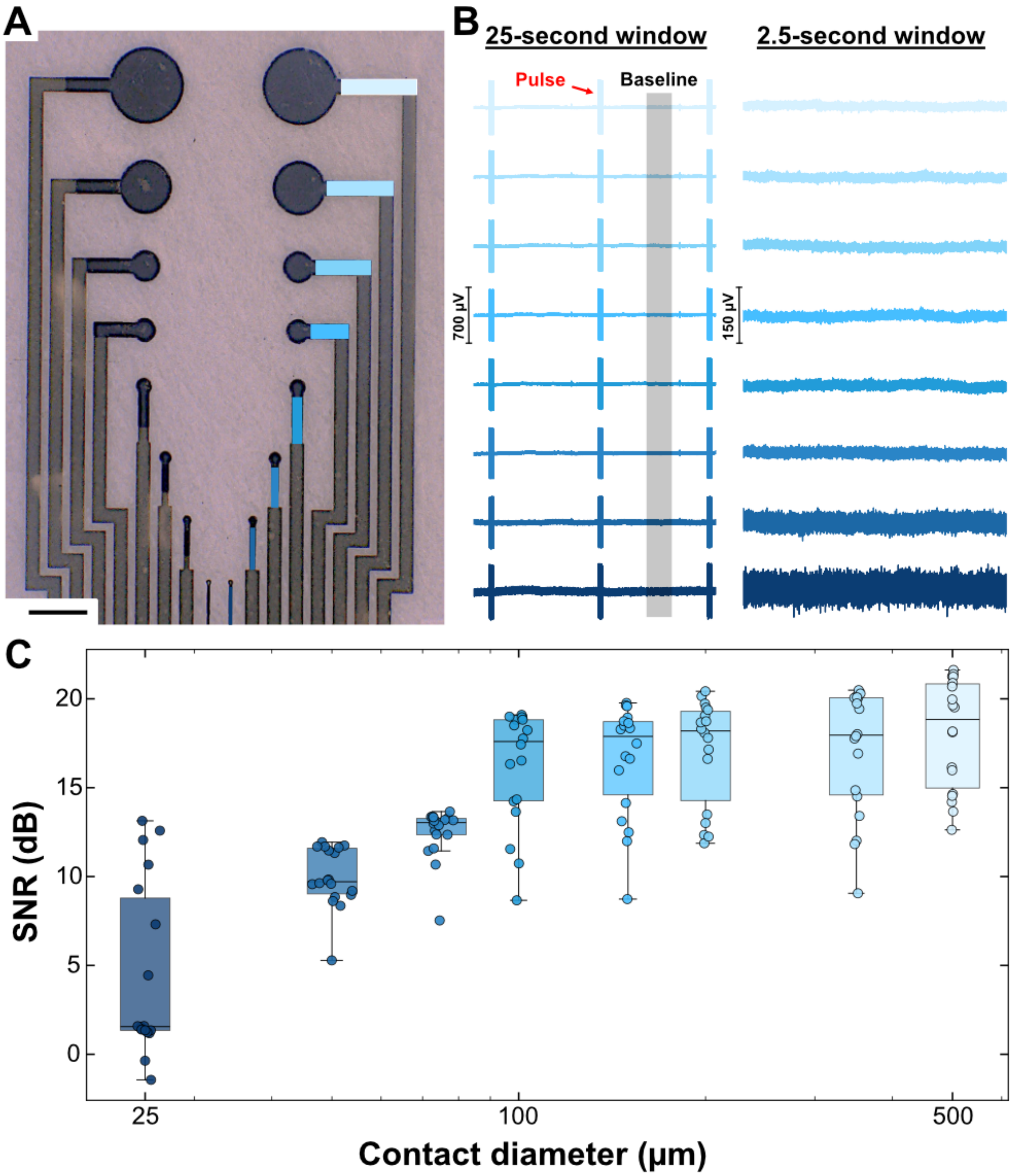


**Figure S8. Relating electrode size to SNR. (A)** Representative image of the $Ti_3C_2T_x$ microelectrode array. Scale bar = 500 µm. **(B)** *Left:* Three representative stimulation pulses from each electrode over a 25-second window, with the color corresponding to the electrode, on the left. *Right:* Zoomed-in 2.5-second clip of noise floor for each electrode diameter. **(C)** SNR as a function of contact diameter. N = 3 devices, 10 stim pulses per recording.

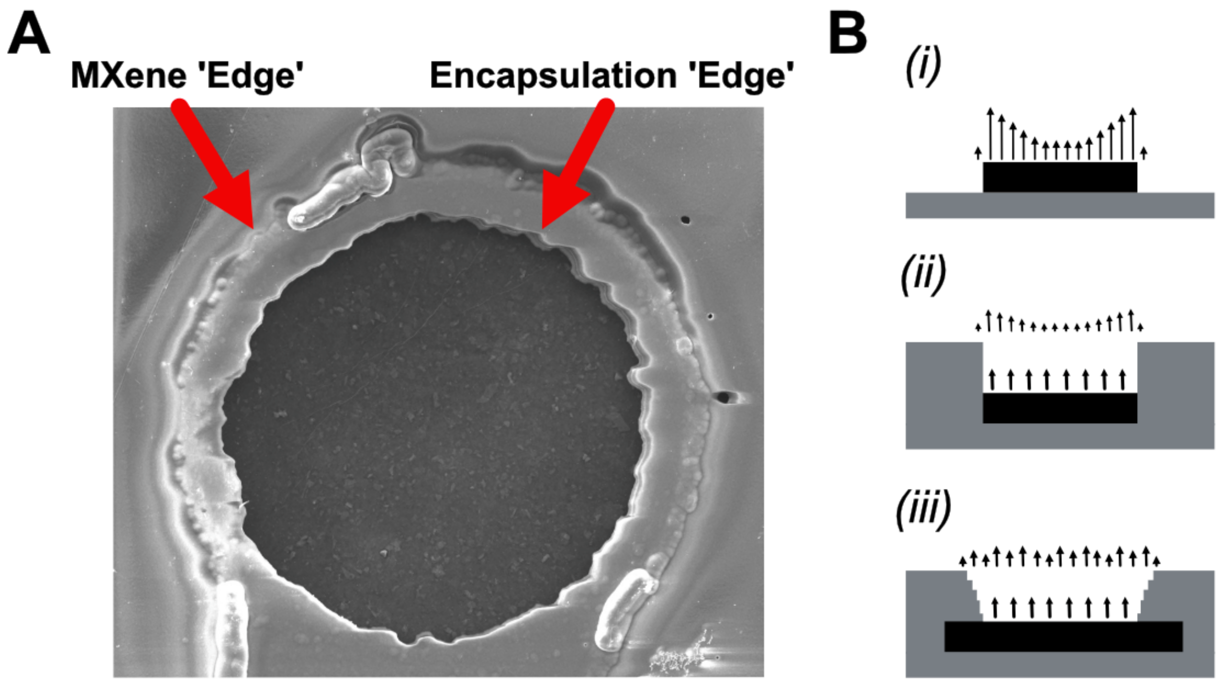


**Figure S9. $Ti_3C_2T_x$ MXene thin-film microelectrode 'edge effects' considerations. (A)** SEM image of a 100 μm diameter electrode contact. Red arrows point to the 'edge' of the spray-coated MXene, defined by the photoresist during fabrication, and the encapsulation edge, defined by the RIE etching of the parylene C encapsulation. **(B)** Expected charge distribution of electrodes fabricated with a *(i)* surface-mounted, *(ii)* simple recessed, and *(iii)* conically / irregularly recessed electrode.[1]

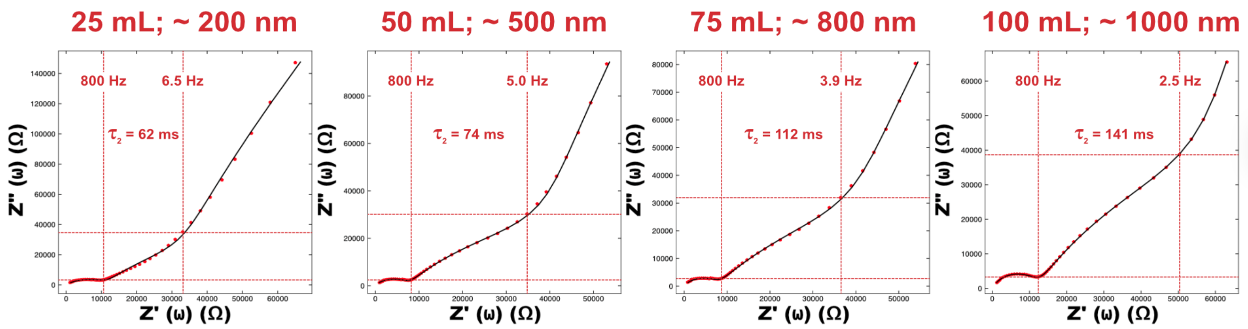


**Figure S10. Emergence of low-frequency RC feature with increasing film thickness.** As the thickness of the spray-coated $Ti_3C_2T_x$ MXene film increases for identically fabricated, 200 μm diameter electrodes, the frequency width and characteristic prominence of the 'bulk-layered' RC time-constant grow. N= 8 electrode samples per film thickness.

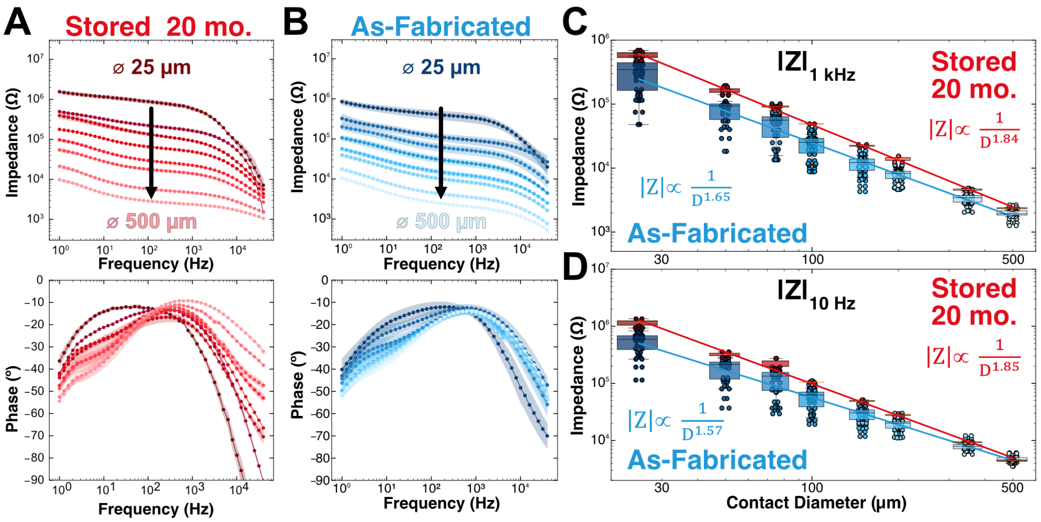


**Figure S11. Shelf-life of $Ti_3C_2T_x$ MXene microelectrodes after 20 months: electrochemical impedance spectroscopy.** Impedance modulus and phase spectra of $Ti_3C_2T_x$ microelectrodes of varying diameter (**A**) stored for 20 months, and (**B**) as-fabricated. Individual points represent the mean, shaded areas represent ± 1 standard deviation. (**C, D**) Impedance modulus at (**C**) 1 kHz and (**D**) 10 Hz as a function of electrode diameter.

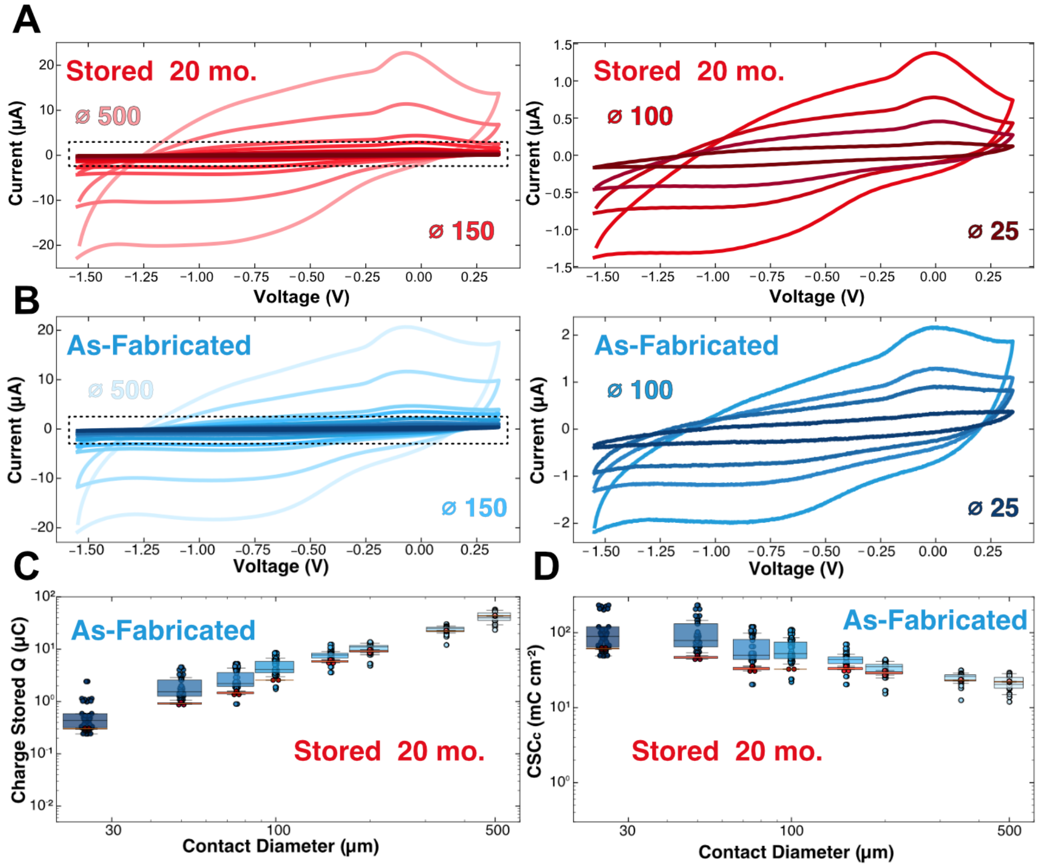


**Figure S12. Shelf-life of $Ti_3C_2T_x$ MXene microelectrodes after 20 months: cyclic voltammetry and charge storage capacity. (A)** *Left:* Cyclic voltammograms averaged across electrodes ($n_{stored}$ = 4 electrodes, each size) of $Ti_3C_2T_x$ MXene microelectrodes stored for over 20 months from -1.55 V (cathodic limit) to 0.35 V (anodic limit). *Right:* Zoomed-in voltammograms for contacts ranging from 100 µm to 25 µm. **(B)** *Left:* Cyclic voltammograms of freshly fabricated $Ti_3C_2T_x$ MXene microelectrodes averaged across electrodes ($n_{as\text{-}fabricated}$ = 34 electrodes, each size) from -1.55 V (cathodic limit) to 0.35 V (anodic limit). *Right:* Zoomed-in voltammograms for contacts ranging from 100 µm to 25 µm. **(C)** Charge stored across all contact sizes for aged and as-fabricated $Ti_3C_2T_x$ devices. **(D)** Cathodic charge storage capacity across all contact sizes of aged and freshly fabricated $Ti_3C_2T_x$ devices.

### SUPPLEMENTARY TABLES

**Table S1. Impedance values at 1 kHz and 10 Hz for $Ti_3C_2T_x$ and Pt devices.**

| Contact Diameter | Frequency (Hz) | $Ti_3C_2T_x$ | | Platinum | |
|---|---|---|---|---|---|
| | | Mean (kΩ) | S.D. (kΩ) | Mean (kΩ) | S.D. (kΩ) |
| 500 µm | 1,000 | 1.97 | 0.34 | 1.11 | 0.11 |
| | 10 | 4.67 | 0.58 | 74.4 | 12.2 |
| 350 µm | 1,000 | 3.40 | 0.67 | 1.92 | 0.20 |
| | 10 | 8.07 | 1.41 | 145 | 19.6 |
| 200 µm | 1,000 | 7.97 | 1.80 | 4.32 | 1.01 |
| | 10 | 21.2 | 5.72 | 347 | 201 |
| 150 µm | 1,000 | 11.6 | 3.57 | 6.70 | 1.77 |
| | 10 | 28.3 | 7.98 | 529 | 277 |
| 100 µm | 1,000 | 23.6 | 8.97 | 13.7 | 4.22 |
| | 10 | 58.2 | 22.7 | 1,100 | 527 |
| 75 µm | 1,000 | 49.8 | 23.6 | 18.5 | 7.13 |
| | 10 | 117 | 50.8 | 1,390 | 755 |
| 50 µm | 1,000 | 90.8 | 42.0 | 49.0 | 17.6 |
| | 10 | 188 | 71.9 | 2,670 | 1,170 |
| 25 µm | 1,000 | 328 | 168 | 140 | 74.7 |
| | 10 | 550 | 218 | 4,370 | 1,960 |

**Table S2. Summary of all equivalent circuit model parameters for 200 µm $Ti_3C_2T_x$ and Pt microelectrodes.**

| Electrode | Circuit Elements | Values from Fitting | Calculated Cdl (µF/cm$^2$) |
|---|---|---|---|
| **$Ti_3C_2T_x$** | $R_s$ (kΩ) | **0.880 ± 0.211** | |
| | $\alpha_0$ | .680 ± .014 | **57.3 ± 4.31** |
| | $Y_0$ (S $s^{\alpha}$) | (6.78 ± .061) x$10^{-8}$ | |
| | $R_{ct}$ (kΩ) | **9.15 ± 0.997** | |
| | $R_{eq}$ (kΩ) | 102 ± 13.7 | |
| | $W_{diff0}$ (kΩ) | 0.276 ± .022 | |
| | $W_{diff1}$ (s) | (2.58 ± .103) x$10^{-4}$ | |
| | $\alpha_1$ | .640 ± .019 | **5,960 ± 982** |
| | $Y_1$ (S $s^{\alpha}$) | (3.63 ± .161) x$10^{-6}$ | |
| **Platinum** | $R_s$ (kΩ) | **0.994 ± 0.153** | |
| | $\alpha_0$ | .814 ± .023 | **66.9 ± 4.87** |
| | $Y_0$ (S $s^{\alpha}$) | (3.39 ± .049) x$10^{-8}$ | |
| | $R_{ct}$ (kΩ) | **1,850 ± 236** | |

**Table S3. Summary of key model parameters for $Ti_3C_2T_x$ and Pt devices from EIS fitting.**

| Contact Diameter | Frequency (Hz) | $Ti_3C_2T_x$ | | Platinum | | Theory |
|---|---|---|---|---|---|---|
| | | Mean (kΩ) | S.D. (kΩ) | Mean (kΩ) | S.D. (kΩ) | |
| 500 µm | $R_s$ (kΩ) | 0.540 | .053 | 0.603 | .055 | 0.720 |
| | $C_{dl}$ (µF/cm$^2$) | 3,820 | 851 | 57.5 | 3.23 | |
| | $R_{ct, surf}$ (kΩ) | 2.29 | .615 | 585 | 83.3 | |
| 350 µm | $R_s$ (kΩ) | 0.710 | .159 | .921 | .185 | 1.03 |
| | $C_{dl}$ (µF/cm$^2$) | 4,500 | 909 | 66.5 | 4.12 | |
| | $R_{ct, surf}$ (kΩ) | 3.93 | .877 | 977 | 182 | |
| 200 µm | $R_s$ (kΩ) | 0.880 | .211 | 0.994 | .153 | 1.80 |
| | $C_{dl}$ (µF/cm$^2$) | 6,020 | 1,031 | 66.9 | 4.87 | |
| | $R_{ct, surf}$ (kΩ) | 9.15 | 0.997 | 1,850 | 236 | |
| 150 µm | $R_s$ (kΩ) | 1.20 | .346 | 1.64 | .349 | 2.40 |
| | $C_{dl}$ (µF/cm$^2$) | 7,830 | 1,440 | 64.7 | 5.38 | |
| | $R_{ct, surf}$ (kΩ) | 13.1 | 1.54 | 2,580 | 610 | |
| 100 µm | $R_s$ (kΩ) | 1.72 | .641 | 2.45 | .682 | 3.60 |
| | $C_{dl}$ (µF/cm$^2$) | 8,470 | 1,970 | 83.0 | 8.94 | |
| | $R_{ct, surf}$ (kΩ) | 25.4 | 6.78 | 4,390 | 855 | |
| 75 µm | $R_s$ (kΩ) | 2.41 | .950 | 3.26 | 1.13 | 4.80 |
| | $C_{dl}$ (µF/cm$^2$) | 8,520 | 2,100 | 102 | 9.03 | |
| | $R_{ct, surf}$ (kΩ) | 49.4 | 8.49 | 4,840 | 970 | |
| 50 µm | $R_s$ (kΩ) | 4.02 | 1.23 | 5.16 | 1.42 | 7.20 |
| | $C_{dl}$ (µF/cm$^2$) | 14,300 | 4,580 | 145 | 11.62 | |
| | $R_{ct, surf}$ (kΩ) | 89.9 | 10.2 | 10,200 | 1,185 | |
| 25 µm | $R_s$ (kΩ) | 8.35 | 1.84 | 6.06 | 1.21 | 14.40 |
| | $C_{dl}$ (µF/cm$^2$) | 27,600 | 7,590 | 213 | 14.72 | |
| | $R_{ct, surf}$ (kΩ) | 381 | 40.9 | 11,200 | 1,460 | |

**Table S4. Charge and cathodic charge storage capacity ($CSC_c$) for $Ti_3C_2T_x$ and Pt microelectrodes.**

| Contact Diameter | $Ti_3C_2T_x$ | | Platinum | |
|---|---|---|---|---|
| | Charge (µC) | CSC (mC/cm$^2$) | Charge (µC) | CSC (mC/cm$^2$) |
| **500 µm** | 40.8 ± 11.4 | 20.8 ± 5.83 | 1.31 ± 0.105 | 0.669 ± 0.054 |
| **350 µm** | 23.9 ± 3.74 | 24.8 ± 3.89 | 0.671 ± 0.056 | 0.698 ± 0.058 |
| **200 µm** | 10.2 ± 2.96 | 32.5 ± 9.44 | 0.247 ± 0.053 | 0.786 ± 0.170 |
| **150 µm** | 7.67 ± 1.68 | 43.4 ± 9.49 | 0.164 ± 0.050 | 0.925 ± 0.281 |
| **100 µm** | 4.69 ± 1.79 | 59.7 ± 22.8 | 0.073 ± 0.008 | 0.936 ± 0.104 |
| **75 µm** | 2.72 ± 1.18 | 61.5 ± 26.6 | 0.045 ± 0.005 | 1.02 ± 0.106 |
| **50 µm** | 1.87 ± 1.06 | 95.2 ± 53.8 | 0.037 ± 0.019 | 1.87 ± 0.955 |
| **25 µm** | 0.588 ± 0.48 | 119.9 ± 97.3 | 0.021 ± 0.015 | 4.29 ± 2.99 |

**Table S5. Capacitances calculated from $C_{dl}$ in the impedance equivalent circuit model and specific capacitance from CV for $Ti_3C_2T_x$ and Pt microelectrodes.**

| Contact Diameter | $Ti_3C_2T_x$ | | Platinum | |
|---|---|---|---|---|
| | Double Layer Capacitance (µF/cm$^2$) | Specific Capacitance (µF/cm$^2$) | Double Layer Capacitance (µF/cm$^2$) | Specific Capacitance (µF/cm$^2$) |
| **500 µm** | 3820 ± 851 | 4060 ± 942 | 57.5 ± 3.23 | 49.2 ± 3.85 |
| **350 µm** | 4500 ± 909 | 4390 ± 892 | 66.5 ± 4.12 | 46.0 ± 4.05 |
| **200 µm** | 6020 ± 1031 | 5550 ± 1180 | 66.9 ± 4.87 | 47.5 ± 3.38 |
| **150 µm** | 7830 ± 1440 | 7590 ± 1240 | 64.7 ± 5.38 | 124 ± 7.69 |
| **100 µm** | 8470 ± 1970 | 8760 ± 1960 | 83.0 ± 8.94 | 113 ± 9.92 |
| **75 µm** | 8520 ± 2100 | 8930 ± 2390 | 102 ± 9.03 | 140 ± 13.4 |
| **50 µm** | 14300 ± 4580 | 13100 ± 4130 | 145 ± 11.62 | 381 ± 53.7 |
| **25 µm** | 27600 ± 7590 | 19300 ± 7640 | 213 ± 14.72 | 249 ± 21.1 |

**Table S6. Comparative electrochemical parameters of $Ti_3C_2T_x$, sputtered Pt, gold, PEDOT: PSS / Pt, Sputtered Pt, SIROF, PtIr, and CNT microelectrodes across the microscale from 500 µm to 18 µm.**[2–5]

| | Diam. (µm) | 1 kHz \|Z\| (kΩ) | 10 Hz \|Z\| (kΩ) | $R_s$ (kΩ) | $CSC_c$ (mC cm$^{-2}$) | $Curr_{max}$ (µA) | $CIC_c$ (µC cm$^{-2}$) | Ref. |
|---|---|---|---|---|---|---|---|---|
| $Ti_3C_2T_x$ | 500 | 1.97 | 4.67 | 0.54 | 20.8 | 1880 | 480 | This Work |
| | 150 | 11.6 | 28.3 | 1.20 | 43.4 | 231 | 654 | |
| | 100 | 23.6 | 58.2 | 1.72 | 59.7 | 97.3 | 619 | |
| | 50 | 90.8 | 188 | 4.02 | 95.2 | 30.6 | 779 | |
| Sputtered Pt | 500 | 1.11 | 74.4 | 0.603 | 0.669 | 134 | 34.0 | This Work |
| | 150 | 6.70 | 529 | 1.64 | 0.925 | 33.6 | 95.0 | |
| | 100 | 13.7 | 1,100 | 2.45 | 0.936 | 22.9 | 146 | |
| | 50 | 49.0 | 2,670 | 5.16 | 1.87 | 5.60 | 144 | |
| Gold | 500 | 4.5[a] | 300[a] | 0.8[a] | 0.1[a,b] | 300[a,c] | 200 [a,b,c] | [2,3] |
| | 150 | 70[a] | 5,000[a] | 2.5[a] | 0.5[a,b] | 30[a,c] | 200 [a,b,c] | |
| | 100 | 170[a] | 10,500[a] | 4.2[a] | 0.8[a,b] | 12[a,c] | 200 [a,b,c] | |
| | 50 | 500[a] | 30,500[a] | 8.5[a] | 1[a,b] | 3[a,c] | 200[a,b,c] | |
| PEDOT: PSS / Pt | 500 | 2[a] | 10.5[a] | 1.5[a] | 3.5[a,b] | 850[a,c] | 650[a,b,c] | [2,3] |
| | 150 | 4.5[a] | 190[a] | 2.5[a] | 5[a,b] | 350[a,c] | 2700[a,b,c] | |
| | 100 | 6[a] | 300[a] | 5.2[a] | 5[a,b] | 150[a,c] | 2700[a,b,c] | |
| | 50 | 15[a] | 1,000[a] | 9.5[a] | 6[a,b] | 40[a,c] | 2700[a,b,c] | |
| Sputtered Pt | 500 | 8[a] | 300[a] | 1.5[a] | 5.1[a,b] | 750[a,c] | 500 [a,b,c] | [2,3] |
| | 150 | 75[a] | 3,000[a] | 2.5[a] | 6.0[a,b] | 100[a,c] | 750 [a,b,c] | |
| | 100 | 200[a] | 5,000[a] | 4.2[a] | 6.2[a,b] | 50[a,c] | 850 [a,b,c] | |
| | 50 | 450[a] | 10,500[a] | 9.5[a] | 8.2[a,b] | 10[a,c] | 850 [a,b,c] | |
| SIROF | 400 | 0.501[a] | 2.51[a] | N/A | 133 | 5,966[d] | 1,900 | [4] |
| PtIr Wire | 18 | 1,100[a] | 60,000[a] | N/A | 1.2 | 42.5[d] | 150 | [5] |
| CNT Fiber | 18 | 200[a] | 4,000[a] | N/A | 372 | 1,570[d] | 6,520 | [5] |

a – Represents values extracted from plots from the respective references

b – Represents a roughly doubled value due to cathodic + anodic normalization

c – Represents values enhanced by positive voltage bias during measurements

d – Represents values calculated from provided pulse-width, CIC, and surface area

**Table S7. Max current and cathodic charge injection capacity of $Ti_3C_2T_x$ microelectrodes at varying pulse width.**

| $Ti_3C_2T_x$ MXene | | | | | | | |
|---|---|---|---|---|---|---|---|
| **Contact Size** | **Parameter** | **100 μs** | **250 μs** | **500 μs** | **750 μs** | **1000 μs** | **1500 μs** |
| **500 μm** | **Current (μA)** | 3882.2 ± 540.4 | 2976.8 ± 321.4 μA | 1884.1 ± 200.1 | 1436.8 ± 151.5 | 1598.8 ± 31.7 | |
| | **CIC (μC/cm²)** | 197.7 ± 27.5 | 379.0 ± 40.9 | 479.7 ± 50.9 | 548.8 ± 57.8 | 814.2 ± 16.1 | |
| **350 μm** | **Current (μA)** | 2504.0 ± 253.3 | 1638.4 ± 200.0 | 891.9 ± 128.7 | 679.2 ± 48.9 | 1629.4 ± 49.1 | |
| | **CIC (μC/cm²)** | 260.2 ± 26.3 | 425.7 ± 52.0 | 463.5 ± 66.9 | 529.5 ± 38.1 | 1693.6 ± 51.0 | |
| **200 μm** | **Current (μA)** | 673.1 ± 86.2 | 545.0 ± 91.6 | 364.9 ± 51.2 | 313.6 ± 51.5 | 266.8 ± 36.1 | 219.2 ± 29.7 |
| | **CIC (μC/cm²)** | 214.2 ± 27.4 | 433.7 ± 72.9 | 580.8 ± 81.5 | 748.7 ± 123.1 | 849.4 ± 115.0 | 1046.9 ± 142.2 |
| **150 μm** | **Current (μA)** | 497.7 ± 2.05 | 368.5 ± 70.8 | 231.1 ± 29.5 | 196.3 ± 19.2 | 177.5 ± 14.6 | 147.5 ± 9.01 |
| | **CIC (μC/cm²)** | 281.6 ± 1.16 | 521.4 ± 100.1 | 653.8 ± 83.4 | 833.5 ± 81.9 | 1004.5 ± 83.1 | 1252.1 ± 76.5 |
| **100 μm** | **Current (μA)** | 282.2 ± 36.5 | 180.0 ± 63.3 | 97.3 ± 15.8 | 103.2 ± 7.3 | 94.7 ± 7.1 | |
| | **CIC (μC/cm²)** | 359.3 ± 46.5 | 573.0 ± 201.6 | 619.4 ± 100.8 | 986.0 ± 70.3 | 1205.9 ± 90.8 | |
| **75 μm** | **Current (μA)** | 186.0 ± 31.9 | 116.3 ± 52.8 | 59.0 ± 18.8 | 48.4 ± 11.3 | 47.3 ± 15.6 | |
| | **CIC (μC/cm²)** | 421.08 ± 72.3 | 658.1 ± 299.1 | 668.1 ± 213.3 | 823.2 ± 192.1 | 1072.8 ± 355.3 | |
| **50 μm** | **Current (μA)** | 60.3 ± 10.7 | 45.9 ± 9.003311 | 30.6 ± 4.18 | 24.9 ± 5.3 | 21.5 ± 2.9 | |
| | **CIC (μC/cm²)** | 307.2 ± 54.9 | 584.6 ± 114.6 | 779.2 ± 106.5 | 952.4 ± 204.6 | 1096.3± 150.0 | |
| **25 μm** | **Current (μA)** | 11.9 ± 4.1 | 18.5 ± 12.1 | 4.88 ± 1.9 | 4.2 ± 2.38 | 5.7 ± 2.8 | |
| | **CIC (μC/cm²)** | 243.8 ± 83.6 | 943.4 ± 616.3 | 497.0 ± 195.5 | 650.2 ± 364.5 | 1172.9 ± 574.2 | |

**Table S8. Max current and cathodic charge injection capacity of sputtered Pt microelectrodes at varying pulse width.**

| Sputtered Platinum | | | | | | | |
|---|---|---|---|---|---|---|---|
| Contact Size | Parameter | 100 µs | 250 µs | 500 µs | 750 µs | 1000 µs | 1500 µs |
| 500 µm | Current (µA) | 623.7 ± 69.7 | 364.3 ± 51.7 | 133.7 ± 8.35 | 426.8 ± 19.2 | 726.7 ± 44.5 | |
| | CIC (µC/cm$^2$) | 31.7 ± 3.5 | 46.3 ± 6.5 | 34.0 ± 2.12 | 163.0 ± 7.3 | 388.7 ± 79.1 | |
| 350 µm | Current (µA) | 472.9 ± 24.8 | 440.6 ± 118.1 | 101.6 ± 3.0 | 397.9 ± 1.88 | 626.8 ± 44.4 | |
| | CIC (µC/cm$^2$) | 49.1 ± 2.58 | 114.4 ± 30.7 | 52.8 ± 1.58 | 310.2 ± 1.47 | 634.2 ± 73.7 | |
| 200 µm | Current (µA) | 122.0 ± 19.5 | 142.1 ± 51.3 | 67.6 ± 15.9 | 74.5 ± 2.59 | 76.5 ± 2.91 | 75.6 ± 2.83 |
| | CIC (µC/cm$^2$) | 38.8 ± 6.21 | 113.1 ± 40.9 | 107.6 ± 25.3 | 178.0 ± 6.20 | 243.6 ± 9.29 | 361.1 ± 13.5 |
| 150 µm | Current (µA) | 89.2 ± 14.1 | 102.1 ± 26.7 | 33.6 ± 2.84 | 74.4 ± 2.75 | 74.3 ± 2.52 | 76.3 ± 3.46 |
| | CIC (µC/cm$^2$) | 50.5 ± 7.98 | 144.4 ± 37.9 | 95.0 ± 8.04 | 315.9 ± 11.7 | 420.6 ± 14.1 | 648.1 ± 11.0 |
| 100 µm | Current (µA) | 37.6 ± 8.8 | 27.488 ± 13.7 | 22.9 ± 8.06 | 21.35 ± 8.09 | 18.0 ± 2.64 | |
| | CIC (µC/cm$^2$) | 47.9 ± 11.2 | 87.3 ± 43.6 | 146.0 ± 51.3 | 203.8 ± 77.3 | 229.8 ± 33.7 | |
| 75 µm | Current (µA) | 29.1 ± 11.6 | 23.6 ± 9.4 | 15.0 ± 4.21 | 14.3 ± 0.28 | 18.2 ± 4.3 | |
| | CIC (µC/cm$^2$) | 65.9 ± 26.3 | 133.6 ± 53.4 | 170.2 ± 47.7 | 243.32 ± 4.8 | 413.0 ± 97.5 | |
| 50 µm | Current (µA) | 29.1 ± 19.6 | 8.3 ± 4.8 | 5.6 ± 2.9 | 4.6 ± 2.4 | 4.3 ± 1.38 | |
| | CIC (µC/cm$^2$) | 148.4 ± 100.2 | 106.1 ± 61.3 | 143.9 ± 74.4 | 176.9 ± 95.2 | 221.5 ± 70.4 | |
| 25 µm | Current (µA) | 4.83 ± 0.93 | 2.76 ± 1.03 | 1.38 ± 0.55 | 1.64 ± 0.61 | 3.11 ± 2.09 | |
| | CIC (µC/cm$^2$) | 98.1 ± 19.0 | 140.7 ± 52.6 | 140.8 ± 56.5 | 250.6 ± 93.4 | 635.0 ± 209.1 | |

**Table S9. Tentative estimates of minimum electrode size based on representative recording and stimulation requirements.** Model equations arising from impedance fitting and CIC values with representative stimulation amplitudes set as magnitude guidelines from other bodies of work.[6–10] Equations from this table are derived from the impedance fit calculation from Figure 2, as well as from the calculation of CIC in Figure 5. Both equations and their corresponding variables are provided below in supplementary equations.

| | **Recording** $D = \left(\frac{\lvert Z\rvert}{e^{b}}\right)^{1/m}$ | | **Stimulation** $D = 2\sqrt{\frac{A_{\min}\,pw}{\pi\,CIC}}$ | | |
|---|---|---|---|---|---|
| **Condition** | Local Fields ($\lvert Z_{10\,Hz}\rvert$ > 1 MΩ) | Multi-Unit Activity ($\lvert Z_{1\,kHz}\rvert$ > 1 MΩ) | Mouse Model $A_{min}$ = 5 µA | Swine Model $A_{min}$ = 50 µA | Human Model $A_{min}$ = 100 µA |
| **MXene** | *m= -1.57, b= 18.19* **D = 16 µm** | *m= -1.65, b= 17.73* **D = 11 µm** | *pw=500, CIC=592* **D = 23 µm** | *pw=500, CIC=592* **D = 73 µm** | *pw=500, CIC=592* **D = 104 µm** |
| **Platinum** | *m= -1.39, b= 19.93* **D = 81 µm** | *m= -1.57, b= 16.69* **D = 6 µm** | *pw= 500, CIC=111* **D = 54 µm** | *pw=500, CIC=111* **D = 169 µm** | *pw=500, CIC=111* **D = 240 µm** |

**Table S10. Influence of processing parameters on the performance of $Ti_3C_2T_x$ MXene microelectrodes.**[11–15]

| Contact Scale | MXene Prep | Diameter (µm) | Area ($mm^2$) | Impedance $\lvert Z_{10Hz}\rvert$ (kΩ) | Area-normalized $\lvert Z_{10Hz}\rvert$ (Ω $mm^2$) | Ref. |
|---|---|---|---|---|---|---|
| **MICRO SCALE** | Thick spray-coat | 75 | 0.00442 | 117 | 516.9 | This Work |
| | Thin spray-coat | 50x50 | 0.00250 | 5,000[a] | 12,500.0 | [11] |
| | Spin-coat | 50x50 | 0.00250 | 500[a] | 1250.0 | [12] |
| | Dip-coat | 100 | 0.00785 | 100[a] | 785.4 | [13] |
| **MACRO SCALE** | Thick spray-coat | 3,000 | 7.06858 | 0.350[a] | 2,474.2 | [14] |
| | Thin spray-coat | 3,000 | 7.06858 | 0.750[a] | 5,301.8 | [14] |
| | Blade-coat | 3,000 | 7.06858 | 0.300[a] | 2,120.7 | [14] |
| | MXtrode | 3,000 | 7.06858 | 0.200[a] | 1,413.8 | [15] |

a – Represents values extracted from plots from the respective references

### SUPPLEMENTARY EQUATIONS

**Equation 1: Diameter scaling with impedance.** Where |Z| is the impedance limit of interest, m, and b are derived from the impedance fit, calculated in Figure 2.

$$D = \left( \frac{|Z|}{e^{b}} \right)^{1/m}$$

**Equation 2: Diameter scaling with charge injection capacity.** Where $A_{min}$ is the minimum current amplitude of interest, *pw* is the pulse-width of the cathodic current pulse, and *CIC* is the charge injection capacity for that material, calculated and provided in Figure 5 for both MXene and Platinum.

$$D = 2\sqrt{\frac{A_{\min} \cdot pw}{\pi CIC}}$$
